# Genetic and epigenetic driven variation in regulatory regions activity contribute to adaptation and evolution under endocrine treatment

**DOI:** 10.1101/2022.02.15.480537

**Authors:** Neil Slaven, Rui Lopes, Eleonora Canale, Diana Ivanoiu, Claudia Pacini, Ines Amorim Monteiro Barbosa, Melusine Bleu, Sara Bravaccini, Sara Ravaioli, Maria Vittoria Dieci, Giancarlo Pruneri, Giorgio G. Galli, Iros Barozzi, Luca Magnani

## Abstract

Comprehensive profiling of hormone-dependent breast cancer (HDBC) has identified hundreds of protein-coding alterations contributing to cancer initiation^1, 2^, but only a handful have been linked to endocrine therapy resistance, potentially contributing to 40% of relapses^1, 3–9^. If other mechanisms underlie the evolution of HDBC under adjuvant therapy is currently unknown. In this work, we employ integrative functional genomics to dissect the contribution of *cis*-regulatory elements (CREs) to cancer evolution by focusing on 12 megabases of non-coding DNA, including clonal enhancers^10^, gene promoters, and boundaries of topologically associating domains^11^. Massive parallel perturbation *in vitro* reveals context-dependent roles for many of these CREs, with a specific impact on dormancy entrance^12, 13^ and endocrine therapy resistance^9^. Profiling of CRE somatic alterations in a unique, longitudinal cohort of patients treated with endocrine therapies identifies non-coding changes involved in therapy resistance. Overall, our data uncover actionable transient transcriptional programs critical for dormant persister cells and unveil new regulatory nodes driving evolutionary trajectories towards disease progression.

## Main

During multicellular development, cell fate is established through a series of heritable transcriptional changes ^14, 15^. These changes are orchestrated by the interaction of transcription factors (TFs) with the regulatory portion of the non-coding genome (*cis*-regulatory elements, CREs) ^16^. CRE activity is largely tissue-specific and contributes to many aspects of cancer aetiology ^17–19^. A large fraction of cancer subtypes displays addiction to the activity of TFs. In line with this, active compounds against nuclear receptors, a targetable class of TFs, account for 16% of the total FDA approved cancer drugs ^20^. Hormone Dependent Breast Cancer (HDBC) cells are strongly dependent on the activity of the nuclear receptor oestrogen receptor (ER*α*), pioneer factors FOXA1 and PBX1 and the transcription factor YY1^10, 16^. These TFs collectively control many cancer hallmarks through their direct interaction with a subset of CREs, particularly distal enhancers ^10, 21–23^. Continuous modulation of ER*α* activity after breast surgery (5 years of adjuvant endocrine therapy) is one the most successful targeted strategies and it represents one of the first examples of precision medicine ^24–27^. Nevertheless, over the course of 20 years post-surgery, cancer returns in up to 50% of patients, suggesting that residual tumour cells can undergo prolonged dormancy ^12, 13, 24^ (Figure 1a).

**Figure 1.**
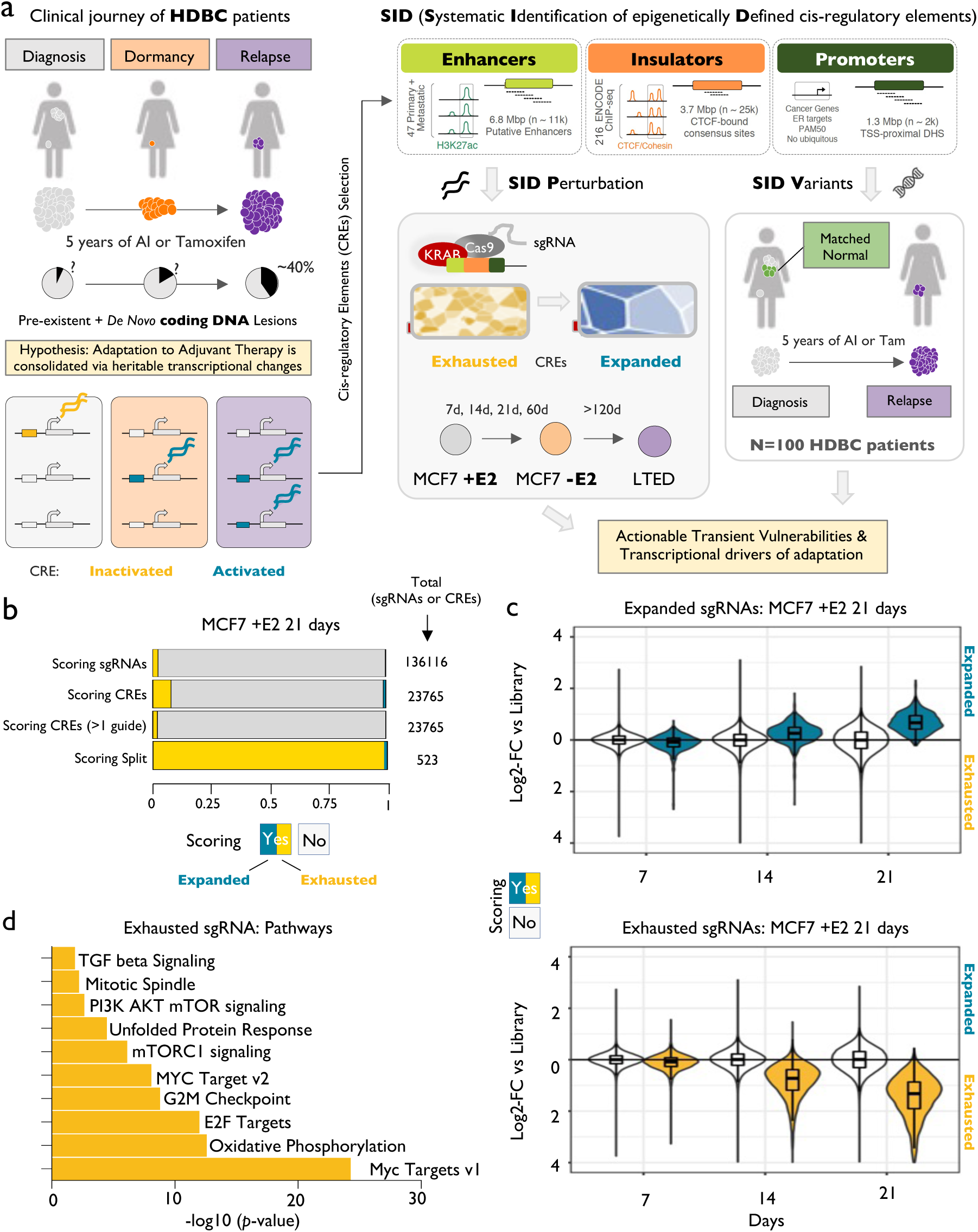
Defining a comprehensive strategy to functionally annotate the non-coding genome of HDBC. **(a)** HDBC journey is characterized by distinct phases. Cells must adapt to different niches and treatments. Overcoming these stresses require profound, heritable transcriptional changes. Leveraging in vivo and in vitro data, we develop SID, a strategy to prioritize HDBC-specific regulatory regions for functional (SID Perturbation) and genomic (SID Variants) annotation in cell line models and patients. **(b)** Bar plot showing the relative fraction of scoring sgRNAs and CREs bearing scoring sgRNAs, upon perturbation of noncoding genome of oestrogen dependent MCF7 cells via SIDP. Scoring sgRNAs showing a significantly decreased frequency at 21 days post-infection are referred to as Exhausted, while those with a significantly higher frequency as Expanded. **(c)** Box plots showing the log2-fold- change of both scoring (either blue or yellow) and non-scoring (white) sgRNAs at 21 days post-infection in oestrogen-dependent MCF7 cells, at 7, 14 and 21 days, as compared to the initial library. **(d)** Bar plot showing the top ten hallmark gene sets enriched among the genes found in the proximity of the CREs with scoring sgRNAs showing a pattern of exhaustion at 21 days post-infection (p-value estimated via a hypergeometric test).

Despite HDBC cells being largely dependent on the activity of these TFs, previous perturbation screens focusing on ER*α* or FOXA1 bound CREs found that only a minority of binding sites appear to be essential for steady-state proliferation *in vitro* ^28, 29^. Yet, TF-centric perturbation has missed CREs driven by additional TFs (*i.e*., YY1 and GATA3 ^30–32)^ and overlooked critical intermediate states in cancer evolution such as adaptive dormancy of persister cells ^12, 13^. To identify CREs contributing to the evolution and adaptation of HDBC tumours exposed to endocrine therapies we developed a prioritised CREs panel (termed Systematic Identification of epigenetically Defined loci, or *SID*) to investigate the role they play both *in vitro* and *in vivo*. The SID panel leverages our patient-derived epigenetic atlas^10^ in which we identified putative enhancers with clonal or sub-clonal representation using Histone 3 Lysine 27 acetylation (H3K27ac) in primary and metastatic HDBC (see Methods). Since disruption of chromatin topology can also contribute to disease evolution in both developmental and cancer models ^33^, SID includes clusters of CTCF binding sites putatively controlling the integrity of topologically associating domain (TAD)^34, 35^ (Figure 1a, Supplementary Figure 1a and Methods).

### Perturbing *SID* regions via CRISPRi

We first investigated the contribution of CREs (at enhancers and TAD boundaries) to HDBC cell growth via massively parallelized dCas9-KRAB (CRISPRi^36^) repressor perturbation. We designed 136,118 single guide RNA (sgRNAs) to interfere with the activity of 23,765 CREs in treatment naïve MCF7 (HDBC cells grown with oestrogen, +E2) (Figure 1a, Supplementary Figure 1b, Supplementary Tables 1 and 2, SID Perturbation or *SIDP*). We reasoned that KRAB-mediated repression mimics CRE loss of function potentially produced by somatic genetic alterations impinging on TF affinity to these sites ^37–39^. *SIDP* covers over 60% of the clonal enhancers active in MCF7 and almost every cluster of CTCF binding sites associated with TAD boundaries (Supplementary Figure 1a).

Nearly 100% of the sgRNAs were captured at high coverage (Supplementary Figure 1b) and then scored based on their relative change after 21 days from infection. This led to the identification of individual sgRNAs either expanded (increased counts corresponding to a potential fitness advantage after the loss of activity of the CRE), exhausted (decreased counts corresponding to a fitness disadvantage after the loss of activity of the CRE) or neutral (Figure 1b). 34% and 0.9% of positive controls and non-targeting sgRNAs scored, respectively, demonstrating the robustness of the approach (FDR <= 0.05; fold-change >= 1.5 or <= -1.5; Supplementary Table 3). Analysis of the temporal dynamics (7, 14 and 21 days) of the sgRNA scoring at 21 days showed reproducible trends (Figure 1c and Supplementary Figure 2d). Interestingly, 98.4% of CREs showing multiple, reproducible scoring sgRNA promote loss of fitness (Figure 1b-c and Supplementary Figure 2d). The regions scoring in our screen showed significant overlaps with observations from previous screens (Supplementary Table 3). Motif analysis on exhausted sgRNAs identified YY1 as the only enriched motif, in line with its critical role in shaping ERα transcriptional activity at clonal enhancers in HDBC ^10^ (Supplementary Figure 2d). Scoring sgRNAs are also associated with many epigenetic features, including KDM5A binding^40, 41^, promoter-specific H3K4me3 and enhancer specific H3K4me1 (Supplementary Figure 2e). Exhausted sgRNAs were significantly associated with CREs near genes controlling metabolic processes (*i.e*., oxidative phosphorylation) and known MCF7 dependencies (MYC targets and PI3K and AKT signalling, Figure 1d and Supplementary Table 3). Collectively, these data establish *SIDP* as a powerful molecular tool for functional characterization of the non-coding genome and demonstrate that only a small fraction of CREs controls cellular proliferation in treatment naïve HDBC cells.

### *SIDP* identifies *de novo* vulnerabilities in adapting cells

Endocrine therapies target disseminated micro-metastatic deposits by interfering with oestrogen receptor activity, reducing the overall chance of relapse by half in patients followed over 20 years ^26, 42^. This effect is largely unpredictable at a single patient level^12, 43^ by virtue of endocrine therapies ability to induce a transient dormant state in persister cells, a process mimicked *in vitro* by long-term oestrogen deprivation^12, 13^. We have shown that *bona fide* coding drivers (*i.e., ESR1* mutations) might not be the actual cause triggering the exit from dormancy as they could emerge and be selected for after awakening, owing to the increased mutational burden associated with replication^12^. We then reasoned that the activity of specific CREs might contribute to the adaptive process occurring during the transition from growth to dormancy entrance^13, 44^.

To investigate this hypothesis, we performed SIDP in long-term oestrogen deprived conditions (-E2), measuring gRNA frequencies at 7, 14, 21 and 60 days after infection (Figure 2a). Analysis of CREs with multiple scoring sgRNAs shows that 10% of these sgRNAs significantly expanded during this period (compared to 1.6% in SIDP +E2, Figure 2b: Supplementary Tables 3 and 4). We interpret this increased representation as a survival advantage emerging uniquely under stress. A significant proportion of sgRNA overlaps between the two conditions and scoring CREs in -E2 were again enriched for YY1 binding motifs, supporting a key role of this TF in the adaptive process, in line with previously reported data ^10^ (Supplementary Figure 4a). In a synergistic lineage tracing study (TRADITIOM, see accompanying manuscript), we show that entrance into dormancy is largely stochastic, with persister dormant lineages selected by chance each time, leading to a significant divergence between replicates ^12^. To test if this process also influences the readout of *SIDP*, we tracked lineages leveraging the non-targeting sgRNAs (n = 501) for up to 60 days of hormone deprivation (full dormancy ^12^). Surprisingly, 210/501 non-targeting sgRNAs (42%, compared to 0.9% in SIDP +E2) showed apparent non-neutral expansion or exhaustion at day 60 (Figure 2c). This behaviour is unpredictable as shown by the evolution of individual non-targeting sgRNA in every replicate (two pools and two replicates, Figure 2b) and by the overall divergent trajectories followed by the two replicates as highlighted by dimensionality reduction (Supplementary Figure 2c). This phenomenon progressively introduces stochastic deviations with time in otherwise predictable perturbation (*i.e.*, ESR1, Figure 2d; SOD1 and CCND1, Supplementary Figure 3a)^28^. These data indicate that the results of a typical CRISPR screen should be taken with care and interpreted in light of these results.

**Figure 2.**
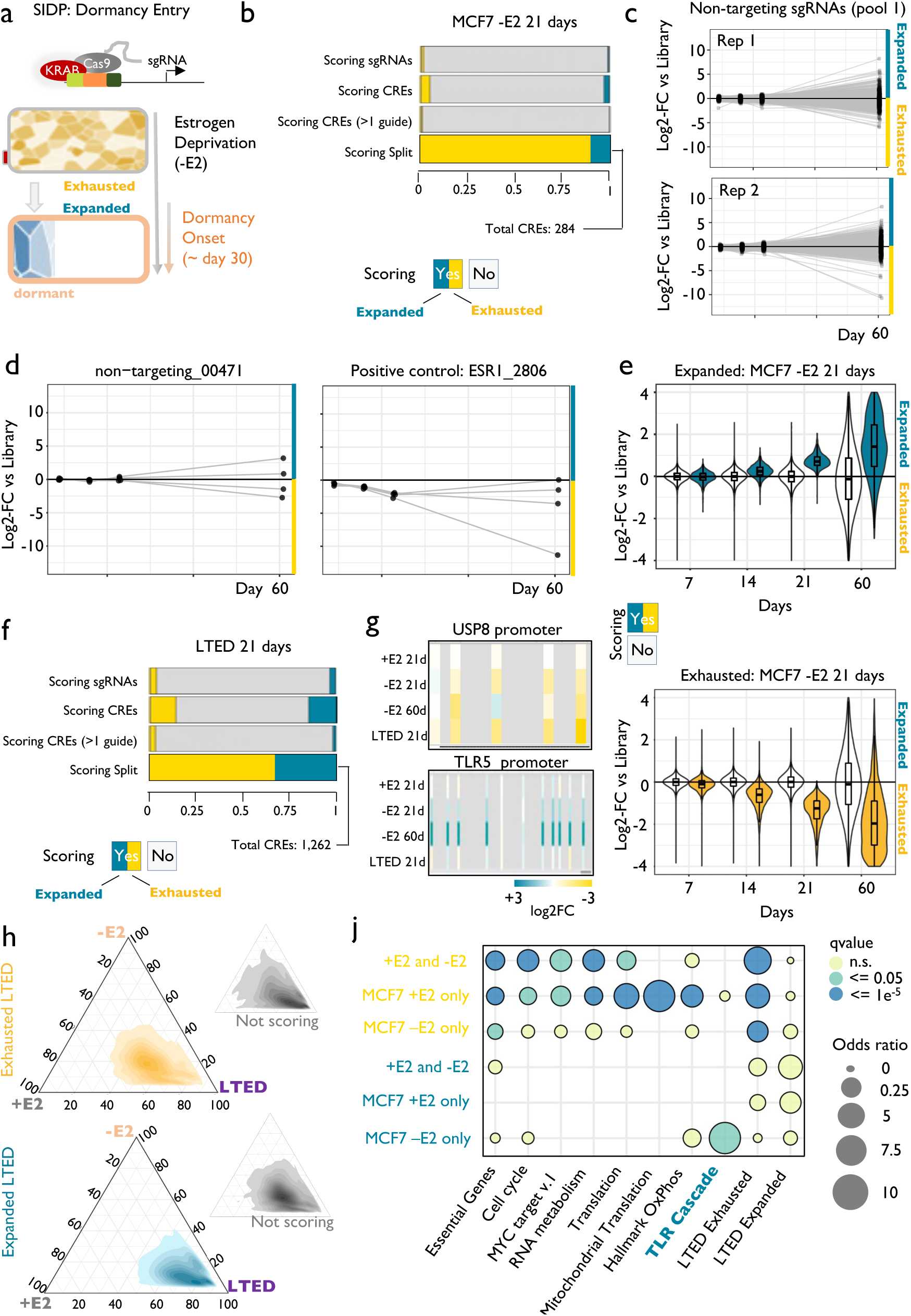
Adaptation to treatment exposes hidden roles for the non-coding genome. **(a)** Experimental design. **(b)** Bar plot showing the relative fraction of scoring sgRNAs and CREs bearing scoring sgRNAs, upon perturbation of noncoding genome of oestrogen-deprived MCF7 cells via SIDP. Scoring sgRNAs showing a significantly decreased frequency at 21 days post-infection are referred to as Exhausted, while those with a significantly higher frequency as Expanded. For the total numbers of sgRNAs and CREs, refer to panel 1b. **(c)** Longitudinal tracking of non-targeting sgRNAs during dormancy entrance (black dots highlight 7-, 14-, 21- and 60-days post-infection). **(d)** Longitudinal tracking of individual non-targeting sgRNAs in four replicates demonstrate stochastic behaviour during dormancy entrance (left panel) as opposed to consistent behaviour of sgRNAs targeting the CRE of essential genes (right panel). **(e)** Box plots showing the log2-fold-change of both scoring (either blue or yellow) and non-scoring (white) sgRNAs at 21 days post-infection in oestrogen-deprived MCF7 cells, at 7, 14 and 21 days, as compared to the initial library. **(f)** Same as panel (b) but for endocrine-therapy resistant cells derived from MCF7 (LTED). **(g)** Summary of the results for the sgRNAs targeting critical CREs of the USP8 and TLR5 genes. **(h)** Ternary plots highlight the higher similarity between LTED and MCF7 +E2 when considering the indicated sets of scoring sgRNAs (Expanded or Exhausted in LTED). **(j)** Bubble plot highlighting the enrichment of distinct biological functions, when considering sets of genes near CREs showing context-specific responses to perturbation.

Nevertheless, our data uncovered a small but significant set of CREs playing a role in the early phases of dormancy entrance (31 CREs with multiple sgRNAs showing a consistent pattern of expansion, Figure 2b). We then systematically compared +E2 and -E2 screens to identify regions showing context-specific behaviour (Supplementary Figure 2d and Supplementary Table 6). During dormancy entrance, MCF7 appear to become independent of several metabolic dependencies, with CREs associated with genes involved in translation, mitochondrial function, and other metabolic processes switching from scoring to non-scoring (+E2>>-E2, Supplementary Figure 2d, e.g., MRPL58 and METTL17, Supplementary Figure 3b). Conversely, a small set of sgRNAs is significantly exhausted exclusively in the -E2 condition, indicating *de novo* vulnerabilities emerging during hormone deprivation (- E2>>+E2, Supplementary Figure 4e-f, e.g., USP8 and SYNV1, Figure 2g and Supplementary Figure 6a). Importantly, the majority of sgRNAs expanding uniquely under therapy showed pronounced enrichment near genes from a single pathway, namely the Toll-receptor activation of the NF-kB pathway (FDR = 0.0049; odds ratio = 13.3; Figures 2e, g, j, Supplementary Figures 4b and Supplementary Table 6). Perturbation of these CREs appeared sufficient to influence the stochastic process controlling dormancy entrance (Supplementary Figures 4c and 5b).

Fully resistant clones emerge from a persister pool after extensive dormancy in both patients and HDBC cell lines models ^12, 45, 46^. Awakening clones exhibit extensive epigenetic reprogramming ^45, 46^ suggesting that the growth of resistant cells might be driven by a distinct set of CREs distinct from that driving the proliferation of the primary tumour. To test this, we run *SIDP* in fully resistant long-term oestrogen deprived (LTED) cells^46, 47^, which represent one fully awakened lineage that emerged from the matched parental MCF7^46, 47^ (Figure. 1a). In line with the results of the screens in +E2 and -E2 MCF7, only a minority of CREs appear to control LTED fitness (Figure 2f; Supplementary Table 5). In stark contrast to proliferating MCF7, the exhausted subgroup does not dominate the scoring sgRNA landscape in LTED (55% vs. 90%, LTED vs. MCF7 +E2), suggesting that LTED have not yet fully adapted. Next, we examined if LTED inherited at least part of the CREs activity acquired during dormancy (Figure 2h). 80% of the dependencies acquired during dormancy appeared to be inherited in LTED (i*.e.,* USP8, Figure 2g-j and Supplementary Figure 7b). Conversely, LTED fitness does not improve upon NF-kB suppression, suggesting that this signalling pathway plays a critical but transient role during dormancy entrance and exit (Figure 2g-j; *i.e*., MYD88 and TLR5, Supplementary Figure 7b). Overall, the application of *SIDP* showed that a relatively small subset of CREs controls different phases of the adaptive process during breast cancer evolution *in vitro*.

### Targeted CRE perturbations accelerate or halt the adaptive processes

*SIDP* demonstrated that cells entering dormancy rapidly switch CREs usage to adapt to treatment (Figure 2 and ^12^). However, the interpretation of the genomic data is difficult due to the stochastic processes influencing individual lineages during dormancy entrance (Figure 2c-d and ^12^). For instance, CREs loss of function conferring fitness advantage under treatment (*i.e.*, TLR/NF-kB) could be explained by three alternative scenarios: increased plasticity (a larger subset of lineages become persister), early awakening and clonal expansion^12^ or complete dormancy bypass (Figure 3a). To test these hypotheses, we tracked the behaviour of cells carrying individual sgRNAs (GFP-NLS) mixed with non-targeting controls during dormancy entrance with live-cell imaging or FACS (Figure 3a).

**Figure 3:**
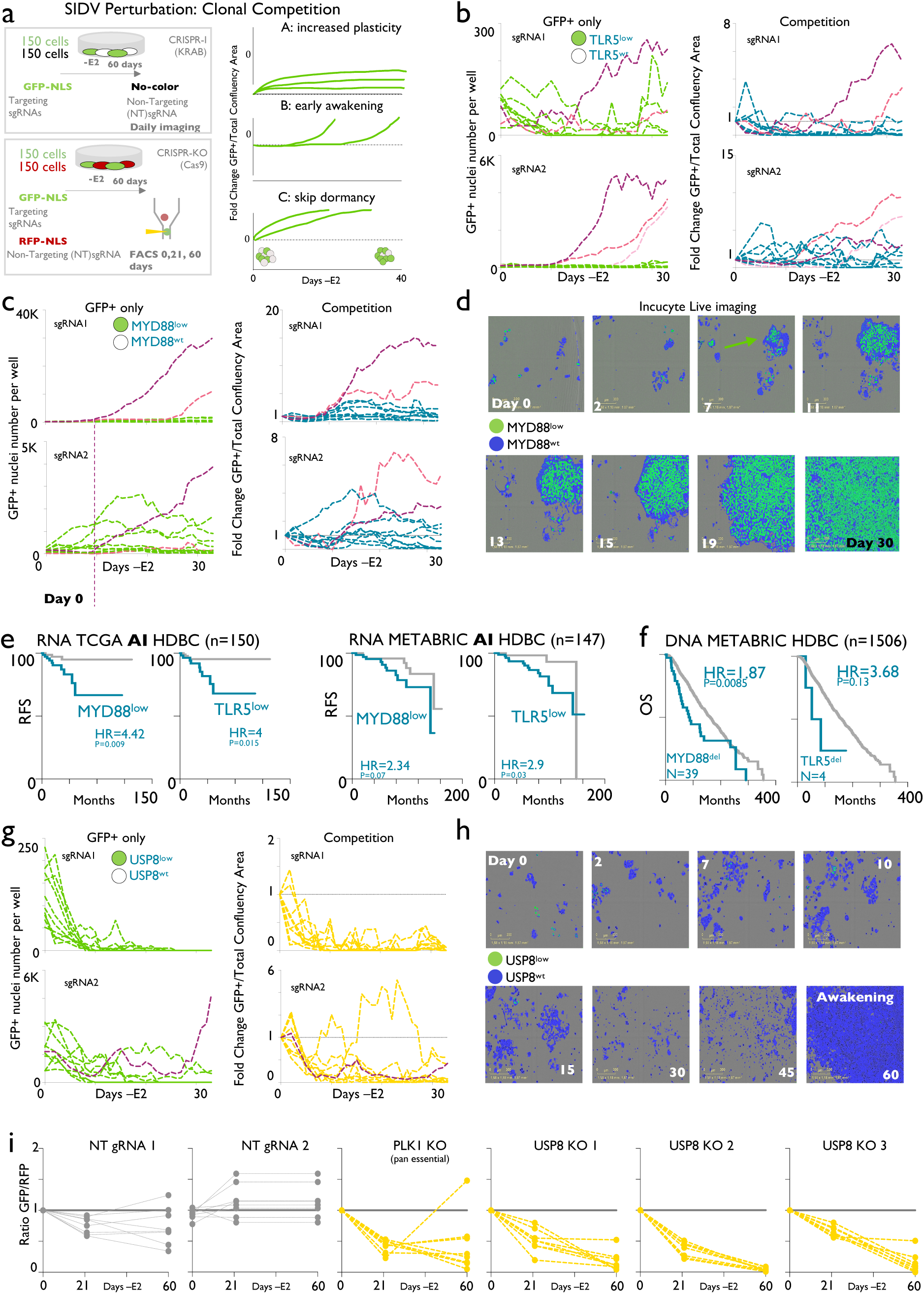
Targeted CRE perturbations accelerate or halt the adaptive processes. **(a)** Overview of the experiments. Cell carrying individual scoring probes were labelled with heritable GFP-NLS are mixed 1:1 with cells carrying non-targeting sgRNA (built-in negative controls). Increased SIDP scores could be explained by three alternative models. **(b-c)** sgRNAs targeting MYD88 and TLR5 accelerate awakening dynamics driving individual clones to early awakening. Green panels: absolute GFP+ count (TLR5 and MYD88 sgRNAs). Blue panels: normalized ratios GFP/non GFP across time points. Pink and purple lines highlight replicates with early awakening events. (**d)** Representative snapshots of the competition between CRISPR-KRAB cells carrying MYD88 targeting sgRNA (green) vs. cells carrying non-targeting sgRNA (blue) throughout dormancy entrance (30 days of continuous estrogen deprivation) **(e-f)** Retrospective patient stratification based on RNA expression or CNVs for MYD88 and TLR5. RFS=recurrence free survival. OS=overall survival. Log-rank p-values calculated with a Mantel-Cox Test. **(g)** sgRNAs targeting USP8 specifically decrease adaptability to oestrogen deprivation. Green panels: absolute GFP+ count (USP8 sgRNAs). Yellow panels: normalized ratios GFP/non GFP across time points. **(h)** Representative snapshots of the competition between CRISPR-KRAB cells carrying USP8 targeting sgRNA (green) vs. cells carrying non-targeting sgRNA 9 (blue) throughout adaptation to estrogen deprivation **(i)** CRISPR-Cas9 knock-out of USP8. FACS sorting was used to quantify green (USP8 sgRNAs carrying cells) and red (non-targeting sgRNAs). FACS analyses were carried out at three specific time points.

To accommodate and quantify the underlying stochasticity of the process, all these experiments were run in ten replicates in absence of cell passaging^12^. Recruitment of KRAB on CREs efficiently led to downregulation of all targets (Supplementary Figure 6a). Cells carrying sgRNAs targeting critical CREs of CCND1 disappear more rapidly in both +E2 and -E2 conditions (Supplementary Figure 6b-c) while MYD88, TLR5 and USP8 targeting sgRNAs do not have any significant impact on the fitness of treatment naïve MCF7 (Supplementary Figure 6b). Conversely, perturbation of MYD88, TLR5 and USP8 gene expression showed a profound effect under oestrogen-deprived conditions. Cells carrying sgRNAs targeting TLR5 or MYD88 showed an accelerated stochastic awakening, with some clones engaging in rapid expansion in days ^12^ (Figure 3b-d). In one case (MYD88 sgRNA #2, pink, Figure 3c), cells showed a behaviour compatible with acquired increased plasticity, given the observed increase in the relative frequency of GFP+ cells in the absence of active cycling. We next stratified independent retrospective cohorts containing only AI-treated patients for MYD88 and TLR5 expression and found that tumours with low pre-treatment expression relapse significantly earlier (HR = 4.42 and 4, *p*-value = 0.009 and 0.015, MTD88 and TLR5 respectively, Log-Rank Mantel-Cox test), in agreement with early awakening (Figure 3f). While MYD88 and TLR5 gene deletions are rare, patients characterized by them also show shorter responses to endocrine treatment (Figure 3f). In summary, these data demonstrate that therapy-induced activation of innate immune signalling plays a central role in entrance and exit from dormancy. In line with this, we find significant evidence that cell-intrinsic activation of this pathway is triggered during active dormancy and suppressed at awakening in single lineages adapting to therapy^12^. Furthermore, cell-intrinsic activation of innate immune signalling is significantly associated with patients with residual disease after neo-adjuvant therapy^48^, suggesting a critical but unexpected association between innate immunity, dormancy and persister cells.

Next, we investigated USP8 as our top *de novo* vulnerability among the *SIDP* hit (Figure 2g and Supplementary Figure 4a). Cells carrying USP8 sgRNA do not have any disadvantage in treatment-naive conditions (Supplementary Figure 9b) while they fail to adapt to -E2 conditions between day 7-30, leading to almost complete eradication (Figure 3g-h). Repeating the long-term competition experiment using a genetic CRISPR-Cas9 system to knock-out the USP8 gene further confirms its vital role in adaptation to endocrine therapies (Fig. 3j). Overall, these data demonstrate that adaptation requires a rapid switch to alternative CREs. Our data show that these emergent phenotypes can be exploited to disrupt or accelerate HDBC cells adaptation to treatment. *In vitro,* these transitions are not the results of Darwinian selection of pre-existent epigenetic clones but are rather induced and become heritable through therapy-induced dormancy ^10, 12, 13^.

### *SIDV* identify patterns of CRE mutations in longitudinal cohorts

*SIDP* is designed to model CRE loss of function via heritable epigenetic repression of CRE activity (KRAB-mediated heterochromatin formation ^49^). Somatic genomic alterations can also strongly influence the activity of individual CREs as well as chromosomal architecture^33, 50^. We reasoned that high-depth genomic sequencing of SID CREs in matched pre-treatment and relapsed samples might shed some insight on the role of the non-coding genome during tumour evolution (Figure 4a). For this purpose, we developed SID variants (*SIDV*, see Methods) and profiled 300 matched samples (normal, primary and relapse biopsies). All patients received either adjuvant Tamoxifen (a selective oestrogen receptor modulator) or Aromatase Inhibitors (Figure 4a and Supplementary Table 7). The median age of diagnosis was 46 for TAM and 58 for AI. Grade and Ki67 status of the primary lesions were similar between cohorts, Figure 4b, Supplementary Figures 7b, e-f and Supplementary Table 7 for the full clinical information). For 58 patients we could also co-profile variants in protein-coding regions, which identified *de novo* drivers of treatment failure (by comparing primary vs. matched relapse) at frequencies comparable to previous studies (i.e. ESR1 mutations^2, 7, 51^, Figure 4c). Using a highly stringent computational pipeline (see Methods and Supplementary Figure 7a), we identified a total of 3576 SNVs and 2,330 INDELs across the cohort, with a median coverage of 117X (Supplementary Table 8). Relapsed samples covered a wide spectrum of anatomic sites and despite showing comparable purity to matched primaries (*p*-value = 0.088), show significantly less genomic alterations (paired two-tailed t-test, *p*-value = 0.0007), potentially indicating decreased genetic intra-tumour heterogeneity due to the bottleneck induced by metastatic seeding (Supplementary Figures 7b-c and 8 a-c). The mutational burden from SIDV regions is highly consistent with previous WGS (Supplementary Figure 7d). Interestingly, the mutational burden is higher in tumours showing high Ki67 and lower in those positive for the progesterone receptor (Supplementary Figure 7e-f). Therapy choice (AI vs TAM) did not seem to impact the number of SNVs at relapse (*p*-value = 0.21; Mann-Whitney Test; Supplementary Figure 8d). We then extended and integrated several machine learning approaches to prioritize the identified 5,524 SNVs and short INDELs based on their predicted effect on TF-binding^52^, chromatin state^53^, accessibility^54^, and splicing^55^ using only models derived from relevant, HDBC-specific genome-wide measurements (Supplementary Figure 7a and Methods). A model-specific *p*-value for each prediction was derived either using permutation-based approaches or by generating a null distribution from the non-coding alterations across all cancer types available in COSMIC ^56^ (see Extended Methods for details).

**Figure 4.**
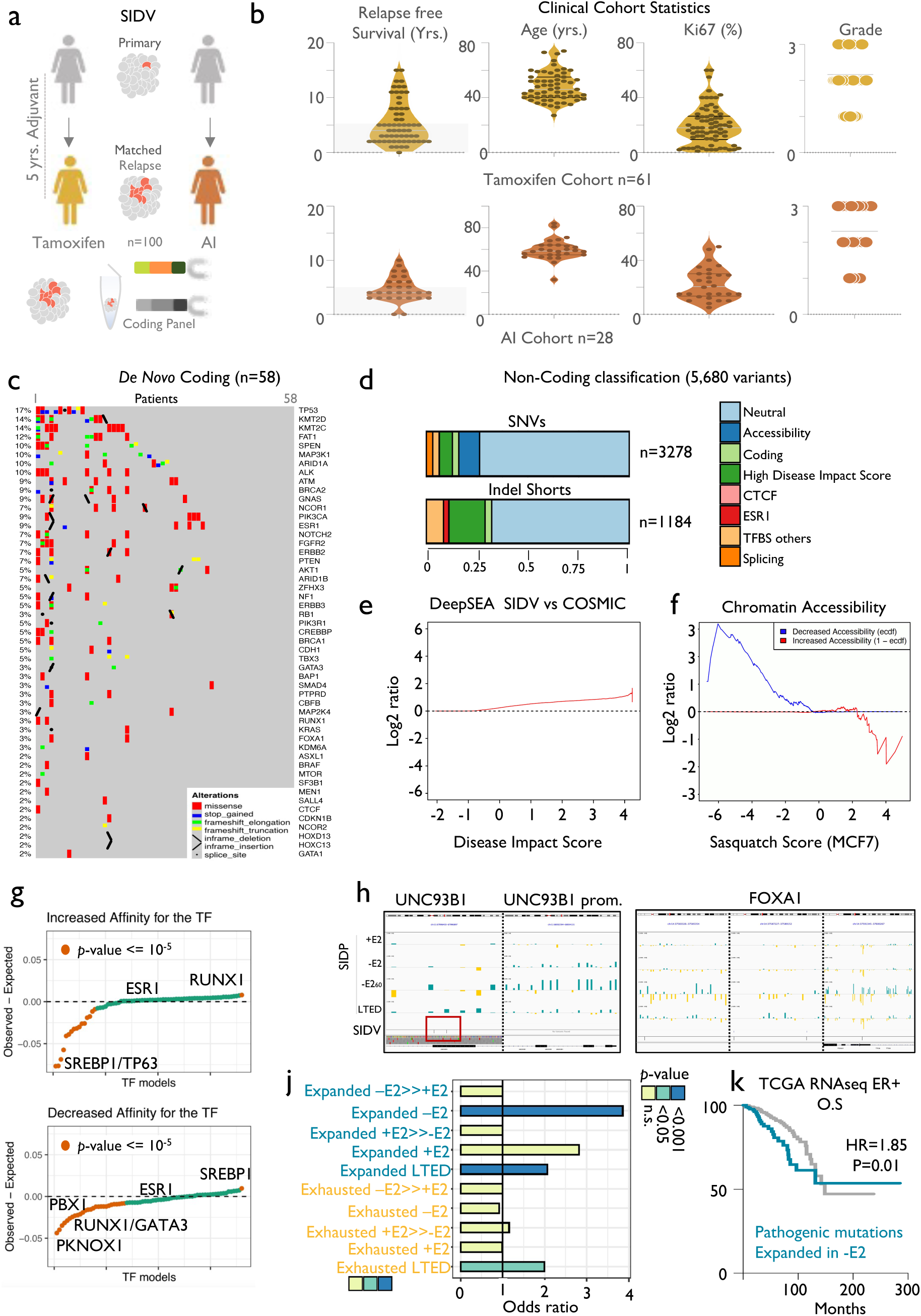
Non-coding variants contribute to heritable transcriptional changes during tumour progression. **(a)** Schematic showing the rationale and implementation of SIDV. **(b)** Overview of the clinical cohorts and the associated features. **(c)** Matched targeted coding profiling identified recurrently mutated (point mutations and indels) genes acquired in metastatic samples. The heat map is showing, for each patient and mutated genes, the type of lesions detected, and the fraction of lesions showing an alteration in each gene (left). **(d)** Pathogenic classification of non-coding variants identified by SIDV. **(e-f)** Functional characterization of SIDV calls as compared to the entire COSMIC catalogue. **(g)** Scatterplot summarising the potential of the profiled SIDV variants to alter transcription factor binding. Each dot represents a TF. TFs are sorted based on their propensity to either increase (top panel) or decrease (lower panel) the affinity to each TF. Values significantly larger than zero indicate a propensity to alter the binding that is higher than expected by chance. Those significantly smaller instead indicate a depletion of variants potentially altering the affinity for a given TF. P-values estimated via Chi-squared Test. **(h)** Integration of SIDV and SIDP identify critical regulators of HDBC biology. SIDP scores and SIDV calls at the indicated loci are shown (IGV genome browser). **(j)** Bar plot showing enrichment of SIDV-identified alterations at sets of regions showing condition-specific patterns upon perturbation (SIDP). P-values estimated via Chi-squared Test. **(k)** Kaplan-Meier plot showing that genes near CREs with an excess of SIDV mutations and overlapping sgRNAs expanded upon oestrogen deprivation (- E2) are associated with prognostic expression levels (HR= 1.85, p-value = 0.01; Log-rank Test).

We predict that ∼up to 30% of SIDV calls might have a functional impact on chromatin (Figure 4d). The Disease Impact Score (as predicted by DeepSEA^57^) of called *SIDV* variants showed significantly higher values than non-coding variants across different cancer types in COSMIC (*p*-value < 1e-16; KS test) (Figure 4e). We also observe enrichment for SNVs with a negative impact on chromatin accessibility (as predicted by Sasquatch^54^; Figure 4f). Variants predicted to exert pathogenic impact on splicing appeared to be under negative selection (our set: 2.28% vs Expected: 4.71%, *p*-value = 9.4e-15, Chi-squared Test). We then focused on those alterations with predicted impact on HDBC-specific TF-binding (as predicted by deltaSVM^52^; see Supplementary Table 13 for the complete information about the TFs considered). Our data show that SNVs potentially altering the binding of several critical HDBC TFs are less frequent than expected (i.e., GATA3, PBX1 Figure 4g and Supplementary Table 12) with the notable exception of SNVs increasing the binding affinity of the HDBC cancer driver RUNX1 or decreasing SREBP1 binding. Interestingly, SNVs with predicted activity (increased or decreased) against ER*α* binding sites do not appear to be under any selective pressure, supporting the notion that most ESR1-bound CREs are not functionally significant^10, 21, 28^. These data suggest that there is an overall negative selection on the binding sites of key TFs. However, when comparing the HDBC- specific alterations we identified to those reported across different cancer types (COSMIC), a residual enrichment for functional alterations was spotted (Figure 4e).

Degeneration and redundancy in the genetic grammar governing cis-regulatory element activity have strongly limited our ability to spot recurrent non-coding mutations^58^. Nevertheless, we hypothesized that by integrating the results from *SIDV* and *SIDP* we could gain more specific insights into the role of non-coding genetic alterations in HDBC (see Extended Methods). Using a lenient threshold (n >= 2; *p*-value <= 0.05; binomial test), 63 *SIDP* CREs showed a significant excess of functional alterations (Supplementary Table 10). These included one CRE falling in a cluster of CTCF binding sites within the UNC93B1 gene, which is part of the genes of the Toll Receptor Cascade whose down-regulation leads to an advantage in -E2 (Figure 2j). Interestingly, both UNC93B1-associated SNVs are predicted to alter splicing while sgRNAs targeting this CRE or UNC93B1 promoter are significantly expanded in either -E2 or LTED screens (but not in +E2 conditions, Figure 4h). Other regions showing both excesses of mutations and *SIDP* significant scores include CREs near FOXA1, a critical TF involved in many aspects of HDBC biology ^21^ (Figure 4h). Furthermore, collapsing the predicted functional mutations at the level of pathways identified an interesting set of biological processes, suggesting that non-coding variants might contribute to promoting cancer evolution by suppressing differentiation and G1 arrest (Supplementary Table 10). Finally, we observed a significant overlap between *SIDV* mutations predicted as potentially pathogenic and *SIDP*, but only when considering CREs bearing expanding sgRNAs under -E2 condition or in LTED cells, suggesting that mutations in these CREs have the potential of conferring a heritable fitness advantage to cells under treatment (Figure 4j and Supplementary Table 10). Mutations found in these CREs tend to show a slight increase in cancer cell fraction in matched metastatic deposits (*p*-value = 0.08; paired samples Wilcoxon Test). Low expression of genes associated with these CREs is associated with poorer prognosis in HDBC (Figure 4k; HR= 1.85, *p*-value = 0.01; Log-rank test). This suggests that cells losing the expression of the target genes due to loss of function of the corresponding CREs might have increased fitness under the selective pressure imposed by endocrine therapies. In support of this, 4/6 of the SNVs in this set show higher cancer cell fraction in matched metastatic samples (*p*-value = 0.03; Chi-squared Test with Yates’ Correction). Taken together, our results demonstrate that nongenetic and genetic mechanisms targeting CREs significantly contribute to tumour evolution by altering the length of therapy-induced dormancy.

## Discussion

The role of the non-coding genome in cancer has been under intense debate ^39, 59, 60^. In this work we have a) established a hormone-dependent breast cancer-specific cistrome^10^; b) systematically perturbed it via targeted epigenetic repression, and c) profiled a large set of somatic alterations accumulated at these regions during tumour evolution. We ran three large-scale perturbation screens against the critical portion of the HDBC non-coding at an unprecedented depth and resolution. We also leveraged a unique patient cohort to profile non-coding genetic alterations longitudinally and at high coverage. Finally, we applied machine learning approaches to systematically dissect the functional consequences of these variants on regulatory potential. Systematic integration of these experimental and computational strategies led to the conclusion that while CREs do not display the strong signature associated with coding drivers, changes in the context-specific regulatory activity of a defined set of CREs plays a crucial role during therapy-induced dormancy. Our results stand out considering the stochastic processes dominating dormancy entrance and exit (see companion manuscript^12^). For example, our *SIDP* screens strongly suggest that signalling converging on NF-kB activation plays a central role in maintaining long-term dormancy. This prediction is corroborated by our transcriptional tracking of single lineages, which shows NF-kB activity being induced in dormant cells but reversed in awakened lineages (see companion manuscript). Of note, mutations on CREs associated with NF-kB regulation are surprisingly infrequent considering the potential benefit to cancer cells under AI pressure (Figure 3g), suggesting that transcriptional switches are the preferred route to adaptation for HDBC cells, possibly because of their reversible nature. In agreement, we could not identify recurrent genetic mechanisms leading to awakening (see companion manuscript). While profiling primary and secondary lesions as an evolutionary endpoint did not reveal many additional therapeutic entry points, transient dormancy might offer an attractive and unexplored stage with potentially actionable transient dependencies. As a proof of concept, we indeed show that targeting USP8 can actively eradicate HDBC once they commit to dormancy. As such, we anticipate that our results will also have critical relevance for the design of future screens that will help expand our knowledge on the regulatory networks underlying therapy-induced dormancy, which we propose as the critical targetable bottleneck in the adaptive journey of breast cancer cells.

## Acknowledgements

All the authors acknowledge and thanks all patients and their families for their support and for donating research samples. The authors gratefully acknowledge infrastructure support provided by Imperial Experimental Cancer Medicine Centre, Cancer Research UK Imperial Centre, National Institute for Health Research (NIHR) Imperial Biomedical Research Centre (BRC) and Imperial College Healthcare NHS Trust Tissue Bank. We thank the NIBR CBT Genomics unit for sequencing support. L.M. was supported by a CRUK fellowship (C46704/A23110). I.B. was supported by CRUK funding (C46704/A23110) and by an Imperial College Research Fellowship. Consent was collected at IEO (European Institute of Oncology, Milan), IOV (Istituto Oncologico Veneto) and IRST (Istituto Tumori della Romagna). Other investigators may have received samples from these same tissues. The views expressed are those of the author(s) and not necessarily those of the NHS, the NIHR or the Department of Health. A special thanks to Xixuan Zhu and Rakshindh Sekhon for their help in the initial crunching of the data, and Giacomo Corleone for help with the initial selection of the SID regions. The authors also thank F. Battiato and A.F. Magnani for their continuous support.

## Contributions

L.M. conceived the original idea. L.M., I.B. and G.G planned and supervised the research. N.S., E.C., carried out the CRISPR validation and *SIDV* assay. R.L., I.A.M., and M.B., carried out the *SIDP* assay. S.B., S.B., M.V.D., and G.P., built the patient cohort. I.B. carried out most of the computational analyses with the help of C.P. D.I. analyzed the coding panel. L.M. wrote the paper with inputs from all authors.

## Material and Methods

### SID panel design

Previous epigenomic annotation of primary and metastatic luminal breast cancer tissues led to the identification of 326,729 putative enhancer regions ^10^. Most of these regions were private or poorly shared amongst individual tumours. However, an overall correlation between the activity of an enhancer in an individual tumour (low ranking index, or RI) and the pervasiveness of its activity across tumours (high sharing index, or SI) was observed. Thus, putative enhancer regions for the panel were biased for those showing a low RI. Starting from the ∼326K regions mentioned above, we first excluded all the private enhancers (RI>=80). 19,482 enhancers were retained and evaluated in terms of their delta of activity between primary and metastatic tumours. The average RI of each enhancer in the primary and metastatic cohorts was calculated (termed RI_Prim and RI_Met, respectively). These two numbers were then used to calculate a region-specific log2(RI_Met/RI_Prim). Putative enhancers showing either higher enrichment in the primary or metastatic samples were selected (regions with RI <=50 in both primary and metastatic, and either in the top positive or negative log2(RI_Met/RI_Prim)). This resulted in 8.05 Mbps covering regions with higher RI in the metastatic samples and 3.7 Mbps showing higher RI in the primary samples. Finally, 2.5 Mbps was assigned to private enhancers being clonal in only 1 or 2 samples. As an internal control, 800 putative enhancer regions were randomly selected among those showing extremely low sharing (SI==1) and ranking (RI==100) index. To reduce the required coverage and to increase the enrichment for potentially functional regulatory regions, DNase-I accessible regions available in ENCODE ^61^ were then used to restrict the area of investigation to the sub-regions within the selected putative regulatory regions. These are more likely to represent clusters of TF-binding sites. To this aim, the regions resulting from the analysis described above were intersected with the DHS from HoneyBadger2 (https://personal.broadinstitute.org/meuleman/reg2map/), which effectively lowered the coverage to ∼9 Mbps. Based on an initial iteration of the capturing strategy, these 9 Mbps were further reduced to about 7, by excluding those regions with either a very low or an extremely high coverage. This resulted into a higher and more even coverage on the majority of the targeted elements Putative insulator regions were selected through a meta-analysis of previously published human ChIP-seq profiles, namely 161 for CTCF (in 89 cell lines or primary cells), 46 for subunits of cohesin (8 targetings SMC3 and 38 targeting RAD21, corresponding to multiple profiles across 5 and 11 cell lines or primary cells, respectively for SMC3 and RAD21) and 8 for ZNF143 (in 4 cell lines or primary cells). ZNF143 has been shown to bind together with CTCF and cohesin and to be specifically enriched at domain boundaries ^62^. Briefly, to identify the strongest, most conserved insulator sites in the human genome, site-specific scoring and spatial clustering of CTCF, cohesin and ZNF143 binding across different cell types were calculated and combined. First, consistently derived, enriched regions from ENCODE datasets ^61^ were downloaded from the UCSC genome browser on July 16^th^, 2016 (Table S1). ChIP-seqs for the same protein in the same cell line (or primary cells) were considered as replicates. Narrow peaks from replicates were merged. The union of the peaks was then computed, and each peak was re-annotated to the sum of the corresponding -log10(p-value) of the overlapping peaks across replicates. To compare the binding profiles across cell types, the obtained scores were converted to percentiles. Given a cell type, percentiles from overlapping CTCF, cohesin and ZNF143 peaks were then summed, resulting in site-specific scores. Separately for each cell type, nearby CTCF-bound regions were then clustered together if found within 10 Kbp from each other. Given each cluster, site-specific scores for each constituent region were combined, first for each cell type, and eventually across all the cell types considered, obtaining an overall score for each cluster. For the final design, the clusters were sorted according to this score, and starting from the highest-scoring cluster, the top clusters covering 3 Mbp of the genome were considered. This way, >95% of previously annotated TAD boundaries^63^ were covered by one or more clusters (keeping in mind the resolution limit of the corresponding HiC datasets, namely 40 Kbp). Promoter regions were selected according to the following strategy. Genes that are either annotated as ER-alpha targets (from the MSigDB Hallmark datasets; PMID: 26771021), found in the PAM50 signature (PMID: 19204204) or being annotated as cancer genes (Network of Cancer Genes version 6.0; PMID: 30606230) while showing an FPKM >= 50 in bulk-RNA-seq data from either LTED, TamR or FulvR resistant cell lines^46^, were considered. From this initial list, genes annotated as housekeeping ^64^were excluded. Promoter regions ([-750, +250] from annotated transcriptional start sites) were derived from the refGene table of the UCSC genome browser on December 13^th^, 2018. Within these regions, only those DNA stretches overlapping DHS (as described above for the putative enhancer regions) were retained. Regions of low mappability along with those mapping to either chromosome Y or the mitochondrial chromosome, as well as those overlapping segmental duplications, were excluded from the design. Regions of unique mappability were defined according to the UCSC genome browser track k50.Unique.Mappability.bb in the Hoffman Mappability collection. After performing an initial, small set of captures, the overall design was further improved by excluding the top and bottom 1% regions. The top 1% regions were responsible for ∼21% of the signal, and the bottom 1% for just ∼0.03% of the signal. Omission of these regions resulted in a more uniform coverage.

### SIDP screens

Two oligo pools for the SIDP library (n=67839 and 69569 oligos respectively, see design information below) were synthesized by Twist Bioscience. Each 60 bp ssDNA oligos contained a 20 bp sgRNA sequence flanked by these sequences 5’-gccatccagaagacttaccg-3’ and 5’-gtttccgtcttcacgactgc-3’ used for PCR amplification and BbsI restriction enzyme-mediated cloning. The oligo pools were cloned into a modified pLKO-TET-ON plasmid by the Golden Gate method and the resulting product was used to transform Endura electrocompetent cells (Lucigen) according to the manufacturer’s protocol. The transformation efficiency was ≈500 fold higher than the SIDP library size and complete and even oligos representation was confirmed by NGS. Large scale preps of bacteria cultures containing the sgRNA plasmid library were harvested using the Genopure plasmid maxi kit (Roche). SIDP library was packaged in lentiviral particles by large scale co-transfection of HEK293T cells with CELLECTA ready-to-use packaging plasmid (Cellecta – cat.no CPCP-K2A) using TRANSIT-LT1 transfection reagent (Mirus biologicals – cat. no. MIR 2300) according to manufacturer guidelines.

MCF7 and LTED cells were engineered to stably express dCas9-KRAB by lentiviral transduction and selected using 10μg/ml blasticidin (Invitrogen) and initially maintained in EMEM (Amimed #1-31S01-I), 10% FBS (Seradigm #1500-500, Lot:077B15), 2mM Glut., 1mM Na Pyr., 10mM HEPES, 1% P/S. Homogeneous dCas9-KRAB expression was confirmed by intracellular staining using Cas9 antibody (Cell Signaling Cat-14697) according to the manufacturer’s protocol.

MCF7-dCas9-KRAB and LTED-dCas9-KRAB cells were then infected with SIDP lentiviral particles at low MOI (≈0.3) in two independent replicates. We transduced ≈1000 cells per plasmid present in the library to guarantee a good representation of all sgRNAs in the population of cells under screening. The cells were selected using 2μg/ml puromycin (Invitrogen) starting at 24 hours post-transduction and maintained in culture in CellStacks (Corning) in the described conditions and for the indicated time points. Cells were then harvested and gDNA isolated using the QIAamp DNA maxi kit (QIAGEN). Amplicons containing the sgRNA sequences were amplified using NEBNext High-Fidelity (NEB) and their representation was analyzed by next-generation sequencing (HiSeq2500, Illumina). During SIDP, for RM condition (full growth media +oestrogen) MCF7-dcas9-KRAB were maintained in DMEM (Gibco #11885-084) supplemented with 10% FBS (Seradigm #1500-500, Lot:077B15), 10mM HEPES, 1mM Sodium-Pyruvate, 1% P/S. For WM (oestrogen-deprived media) MCF7-dcas9-KRAB and LTED were maintained in Phenol-free DMEM (Gibco #11880-28) supplemented with 10% Fetal Bovine Serum, charcoal-stripped, USDA-approved regions (Gibco #12676029), 2mM L-Glutamine, 10mM HEPES, 1mM Sodium-Pyruvate, 1% P/S.

### Flow cytometry-based cell competition assays

MCF7-dcas9KRAB were infected with a modified pLKO-TET-ON lentiviral vector to deliver constitutively expressed sgRNAs in the target cells. Cells transduced with targeting sgRNAs (expressing mCherry) or non-targeting sgRNAs (expressing GFP) were mixed (ratio 2:1 mCherry: GFP) and maintained in culture as described above. At each time point, cells were harvested and analyzed by flow cytometry using CitoFLEX S (Beckman Coulter). We recorded at a minimum of 2,000 single-cells for each condition and the results were analyzed by FlowJo.

### Incucyte-based competition assays

MCF7-dcas9-KRAB cells were engineered by lentiviral transduction containing a vector expressing NLS-eGFP (kindly provided by Dr Chun Fui Lai, Imperial College London). Transduction efficiency was evaluated with EVOS XL Core Imaging System microscope (Thermo Fisher – AMEX100), and a population of bright GFP-positive cells was obtained by Fluorescence-Activated Cell Sorting (FACS). Sorting was performed by the Flow Cytometry facility at MRC London Institute of Medical Sciences. MCF7-NLS-eGFP-dCAS9KRAB were then transduced with lentiviral particles containing plasmids expressing individual sgRNAs and selected with Puromycin (Sigma-Aldrich cat no. P8833). For each gene of interest, 150 eGFP positive (targeting sgRNA) and 150 transparent (NTC-sgRNA) MCF7-dcas-9KRAB cells were seeded per well in a 96 wells ImageLock plate (Sartorius – cat no 4379) both in the presence and absence of oestradiol (Complete medium with 10% FCS +/- 17-ß Oestradiol 1x10-8 M (Sigma Aldrich – cat no E-060)) in parallel, for a total of ten replicates per condition. The plate was routinely media changed and imaged daily with Incucyte (Incucyte ZOOM - Sartorius) using a Dual Color 10X 1.22um/pixel Nikon Air Objective (Sartorius cat no 4464). (Green filter: Ex 440/480 nm, Em 504/544nm). The IncuCyte ZOOM Live-cell analysis system software was used to perform automated cell imaging over time and to calculate cell-by-cell segmentation employing a manually adjusted segmentation mask used to train the images taken at each time point. The total percentage of confluency and the total GFP positive area percentage were automatically registered by the software and used to calculate the ratio between the two parameters normalized to day 0, to highlight an increase (> 1: fitness) or a decrease (< 1:vulnerability) in the trend of GFP-targeting representation over the non-targeting one. Numbers of green nuclei were also automatically counted by the software to obtain the GFP+ only cell count.

### qPCR analysis

RNA was extracted from dcas9-KRAB-MCF7 cells transduced with targeting and non-targeting sgRNA (Qiagen, cat no. 74016). RNA was retrotranscribed using iScript (BioRad, cat no. 1708891). Quantitative PCR was performed with QuantStudio3 Real-Time PCR instrument (Applied Biosystems, cat.no A28567) using an SYBR-green PCR master mix reporter (Applied Biosystems, cat no. 4309155) and the following primers, designed around the promoter of the repressed genes. USP8 fwd: GGGTCTTGGGCCCTAGCA, rvrs: CAGAGCTTGTCTCCGGGGTA - MYD88 fwd:CTGCTCTCAACATGCGAGTG,rvs: CAGTTGCCGGATCTCCAAGT – TLR5 fwd: GCGCGAGTTGGACATAGACT, rvrs: GAGGTTTTCAGGAGCCCGAG).

#### Tissue Specimens

Longitudinal Formalin-Fixed Paraffin-Embedded (FFPE) HDBC samples were retrospectively collected from 100 patients. 61 patients were collected from Professor Giancarlo Pruneri at The European Institute for Oncology, Milan. Samples from 26 patients were collected from Professor Andrea Rocca at The Cancer Institute of Romagna, Meldola. The remaining 14 patient samples were collected from Professor Maria Vittoria Dieci at The Institute of Oncology Padova. The material was collected in the form of 10 µm slices. Detailed clinical notes were provided for each patient including age at diagnosis, Tumour grade, Percentage of ER-positive cells, Percentage of PR positive cells, Percentage of Ki-67 high cells, Percentage of HER2 positive cells, Years until relapse, Metastatic site, Type of Chemotherapy, Type of hormonal therapy. A full summary of the clinical data can be found in Supplementary material 3.

#### Sample Preparation Workflow Extraction

DNA was extracted from 10 micro-meter slices using the Qiagen GeneRead DNA FFPE extraction kit (Qiagen, Catalogue no. 180134) which includes a Uracil N Glycosylase enzyme treatment to reduce FFPE artefacts. DNA quality and quantity were assessed using an Agilent Tapestation 2200 using the Genomic DNA screentape and reagents (Agilent, Catalogue no. 5067-5365 and 5067-5366). Samples were sonicated custom number of cycles to achieve fragments of uniform length. Post-sonication samples were quality controlled using the Tapestation 2200 instrument with a threshold set for samples to have at least 60% of fragments between 100-500bp to proceed with processing. DNA underwent a second treatment with NEBNext FFPE DNA Repair Mix (NEB, Catalogue no. M6630) to further reduce FFPE artefacts.

#### Library Preparation and capture

DNA libraries were prepared from 30 ng – 1 ug of DNA using the NEBNext Ultra 2 DNA library kit for Illumina sequencing. Unique dual 8bp indexes were used for each sample (A gift from Paolo Piazza of the Imperial British Research Council Genomics Facility). DNA libraries from 15 samples were pooled and captured with the SID-V capture probes produced by Twist Biosciences (ratio of 1.5 ug DNA libraries, 100 ng each, to 800 ng of capture probes). Non-captured DNA was recovered using SPRI size selection beads to be used for a secondary capture. Post-capture amplification was performed using the KAPA HiFi Hot Start PCR ReadyMix Kit (KAPA Biosystems, Catalogue no. KK2601). Post-capture amplified libraries were quality controlled and quantified using a Tapestation 2200 with the High Sensitivity reagents.

#### Sequencing

The initial 40 patients were sequenced on an Illumina HiSeq 4000 Instrument (Standard mode, 2 x 150bp). After sequencing the initial 40 patients, sequencing was then performed by Novogene on an Illumina NovaSeq 6000 using 2 x 150bp chemistry. An average of 176 million reads per sample was achieved.

#### Raw data processing of the captured DNA

First, paired-end reads from each sample were trimmed for adapter sequences and based on quality using Trim-galore (version 0.6.4; http://www.bioinformatics.babraham.ac.uk/projects/trim_galore/) in --paired mode. Alignment to the hg38 genome was then performed using bwa mem (version 0.7.15; https://arxiv.org/abs/1303.3997) using default parameters. The hg38 reference genome along with the corresponding annotation and known variant files mentioned in this and the following paragraphs were part of the Broad Institute Bundle, as per download from the Broad FTP on February 5^th^, 2018. Sambamba (version 0.7.1; PMID: 25697820) was then used to convert the resulting SAM to a BAM file (using sambamba view -S -h -F “not unmapped” -f bam). Sambamba sort and index were then used for sorting and indexing the resulting BAM file. The markdup function from Sambamba was used to mark potential PCR duplicates. Recalibration of base quality scores was performed using GATK4 (version 4.1.3.0; ^65^). The BaseRecalibrator function was run (providing dbSNP version 146 via the parameter --known-sites) followed by ApplyBQSR. The resulting BAM file with recalibrated scores was indexed using Sambamba. Final metrics for each sample were computed using the CollectHsMetrics function of the Picard tools (version 2.20.6; http://broadinstitute.github.io/picard/).

#### Mutational calling pipeline

To robustly identify SNVs and short INDELs, a pipeline deriving a consensus between three independent tools (Mutect2, Platypus and Strelka) was deployed. Mutect2 (part of GATK4 version 4.1.3.0;^66^) was run individually on each primary and metastatic sample using the matched normal as reference. The -L option was used to specify the targeted regions. The file af-only-gnomad.hg38.vcf.gz acted as the source of germline variants with estimated allele frequency (as specified via the --germline-resource option). Parameters --af-of-alleles-not-in-resource 0.001, --disable-read-filter MateOnSameContigOrNoMappedMateReadFilter and --f1r2-tar-gz were also specified. The output from running the --f1r2-tar-gz option was then used to learn an orientation biased model (separately for each sample), leveraging the LearnReadOrientationModel function of GATK4. This allows estimating the substitution errors occurring as a result of damage induced by FFPE, by identifying residues showing a significant bias of substitutions on a single strand. The resulting model was then fed into the FilterMutectCalls function of GATK4 so that potentially affected residues can be flagged for subsequent filtering (see below).

Platypus (version 0.8.1.2;^67^) was run on each patient, jointly considering the normal as well the primary and metastatic profiles. The union of the variants called by Mutect2 separately on the primary and metastatic sample (see above) was used as prior (-- source option). Option --minReads was set to 4.

Strelka (version 2.9.10; ^68^) was run independently for each primary and metastatic sample using the matched normal as a reference, with default parameters. While both Mutect2 and Platypus jointly identify SNVs and INDELs, Strelka relies on Manta (version 1.6.0; ^69^) for the detection of INDELs. Manta was run first, and the resulting list of candidate INDELs was then provided to Strelka via the --indelCandidates option. Considering the resulting lists of SNVs and INDELs, both common and tool-specific filters were applied to the lists generated by the different tools. General filters included:

- A minimum depth of 20 reads was applied to both normal and tumour samples.
- A minimum alternate allele coverage of 2 reads.
- Exclusion of variant overlapping known SNPs (dbSNP version 146). Tool-specific filters were set as follows:
- Mutect2: after running FilterMutectCalls (GATK4) which also considered FFPE artefacts as estimated by the orientation bias model, only those variants marked as PASS were retained.
- Platypus: all variants flagged by the tool were discarded, except those marked as PASS or including just one or more of the following flags: badReads, HapScore, alleleBias.
- Strelka: only variants marked as PASS were kept for further analyses.
- Of the resulting filtered variants, only those SNVs or short INDELs that were consistently identified by at least 2 out of 3 calling algorithms, very retained for further investigation.

#### Copy number calling pipeline

CNVkit (version 0.9.7; ^70^) was run in batch mode on the tumour bam files, using all normal bam files of each capturing-sequencing batch as input for the option --normal. SIDV3 intervals were specified under option --targets. The reference genome used for mutational calling was employed (Broad Bundle).

#### Purity and Cancer Cell Fraction estimation

To estimate the Cancer Cell Fraction (CCF) of each SNV, only SNVs with an estimated copy number of 2 were considered. Separately for each sample, the SNVs fulfilling this criterion were hierarchically clustered based on their VAF (using Euclidean distance and complete linkage). The dendrogram was then cut at a fixed height of 0.15, and the cluster with the larger mean VAF was identified. This mean VAF was then used to estimate the purity of the sample: purity = VAF_mean_ * 2. The CCF of each variant was then calculated starting from its VAF and the estimated purity for the sample, using the following formula: CCF = VAF * (2 * (1 - purity) + CNA_TOT * purity) / (CNA_MUT * purity) ^71^. While CNA_TOT was known (2, see above), each variant was assumed to be heterozygous, with CNA_MUT set to be 1 ^71^.

#### Data collection and pre-processing to train the deltaSVM models

A manually curated list of previously published, high-quality human ChIP-seq datasets from luminal breast cancer cell lines was compiled. Only those having a high-quality model (position weight matrix or PWM) describing their binding preferences were considered. The reason behind this choice is that knowing the binding preferences was a prerequisite to generate well-controlled negative sets for the deltaSVM models. Briefly, each PWM was used for genome-wide predictions of binding sites specific for each TF, to then derive a positive (predicted TF-binding site showing a ChIP-seq peak) and a negative (predicted TF-binding site, that could be in principle be contacted by the TF, but without a ChIP-seq peak) training set. This selection resulted in 72 ChIP-seq, corresponding to 43 transcription factors (Table S2). Peaks in BED format were downloaded from the Gene Expression Omnibus (GEO;^72^). Regions in hg18 or hg19 coordinates were converted to hg38 using liftOver^73^, and then filtered against the ENCODE blacklists^74^ using BEDTools ^75^.

#### Predicting the functional effects of the identified variant

Available, pre-computed genome-wide predictions were used to assess the impact of somatic variants on chromatin accessibility (Sasquatch;^54^), mRNA splicing (Splicing Clinically Applicable Pathogenicity prediction or S-CAP;^55^) and protein-coding sequence (Cancer Genome Interpreter or CGI;^76^). Available models based on deep learning (DeepSEA;^57^) were used to compute the overall disease impact score of each variant. Support vector machines (SVMs) were instead trained to predict the impact of somatic variants on the binding affinity of luminal breast cancer-relevant TFs. For each one of the different functional categories, the predictions were obtained as follows:

- Chromatin Accessibility: The Sasquatch R package version 0.1 (https://github.com/Hughes-Genome-Group/sasquatch) was used to assess the impact of the identified somatic variants using the available model pre-trained with *ENCODE_DUKE_MCF7_merged* DNase-seq dataset. Briefly, hg38 coordinates were converted to hg19 using liftOver ^73^. Analysis of multiple reference-alternative alleles pairs was then performed using the *RefVarBatch* wrapper, using *DNase* as fragmentation type: (frag. type = “DNase”) and *human* as propensity source (pnorm.tag = “h_ery_1”). Empirical *p*-values were estimated separately for observing a predicted increase or decrease in accessibility. A *null* distribution was derived from the COSMIC non-coding database ^56^, which contains millions of variants from different cancer types. Version 92 (08.2020) was downloaded as a flat file on October 12^th^, 2020. Sasquatch was run on the entire set of variants, but only those overlapping with the SIDV3 intervals were retained to compute the *null*.
- mRNA splicing: Full S-CAP predictions (scap_COMBINED_v1.0.vcf) were downloaded from http://bejerano.stanford.edu/scap/ on August 27^th^, 2019. A custom Python script was prepared to annotate the somatic variants with these predictions.
- Protein-coding sequence: The list of candidate somatic mutations was submitted to the CGI webserver on December 1^st^, 2020 (https://www.cancergenomeinterpreter.org/). Also, in this case, hg38 coordinates were converted to hg19 using liftOver ^73^.
- Disease impact score: models from DeepSEA version 3 were used to estimate this. Hg38 coordinates were converted to hg19 using liftOver^73^ and a corresponding *null* distribution leveraging COSMIC was computed as described above for chromatin accessibility.
- TF-binding affinity: deltaSVM^52^ was used to predict significant effects of a somatic variant in decreasing on increasing the affinity of the region for a given TF. First of all, for each considered PWM (Table S2) a genome-wide map of the high-affinity sites in the human genome (hg38) was predicted using FIMO ^77^. FIMO was run with the following parameters: --thresh 1e-4 --no-qvalue -- max-stored-scores 10000000, separately for each motif. Regions of unique mappability (as defined according to the UCSC genome browser track k50.Unique.Mappability.bb in the hoffmanMappability collection) were defined using BEDTools^75^, and only those were retained for the next steps. This information was coupled to the corresponding TF-ChIP-seq, to derive a positive (predicted TF-binding site showing a ChIP-seq peak) and a negative (predicted TF-binding site, that could be in principle be contacted by the TF, but without a ChIP-seq peak) training set. Each region in these two sets was defined as the 100 bps of genomic DNA centred on the predicted, high-affinity site. The actual training set used were randomly subsampled versions of these two sets (n = 10,000). Training of the support vector machine (SVM) discriminating the positive from the negative examples was performed by running gkmsvm_kernel (with option -d set to 3) followed by gkmsvm_train. After that, gkmsvm_classify was used to generate a weighted list of all possible 10-mers, where each 10- mer is assigned a SVM weight corresponding to its contribution to the prediction. With this list of weights, it was possible to predict (using the script deltasvm.pl) the impact of any sequence variant on the regulatory activity of a given region. One limitation of this approach when comparing models generated with very different data (like in this case for different TFs) is to define model-specific thresholds. To overcome this, the set of genomic regions under investigation was randomly mutagenized, resulting in a dataset in which every sequence was mutagenized at 5 residues (to all the three possible variants). The resulting values were used to compute model-specific *null* distributions, that were used to estimate empirical *p*-values for the predicted effects of the real set of mutations.

#### Variant classification

A variant was classified as potentially pathogenic if meeting at least one of the following conditions:

- Annotated as either Missense, Nonsense, or Frameshift by the CGI;
- Showing an empirical *p*-value equal or lower than 0.05 in terms of either disease impact score (DeepSEA), or predicted increase or decrease in chromatin accessibility (Sasquatch), or for the affinity of any of the 43 transcription factors considered in the deltaSVM models;
- Showing any of the following S-CAP scores: 1) score >= 0.006 in case of mutations in the introns upstream of a 3’ SS or downstream of a 5’ SS; 2) score >= 0.033 in case of a mutation in the 3’ AG (3’ SS core); 3) score >= 0.009 in case of synonymous exonic mutation; 4) score >= 0.034 for a mutation in the 5’ GT (5’ SS core); 5) score >= 0.005 in case of variants lying in the canonical U1 snRNA-binding site, excluding the 5’ SS core (5’ extended); 6) score >= 0. 006.

#### Identification of regions showing an excess of regulatory mutations in the tumour samples cohort

Given a regulatory element targeted by the enrichment strategy, the probability of a given region to show an excess of mutations predicted as pathogenic was evaluated based on a binomial distribution. The expected probability *p* was estimated as the fraction of variants predicted as pathogenic in the entire datasets. The *pbinom* function from R was used to calculate the probability of seeing an equal or better number of *q* pathogenic variants in the region, given the expected probability *p* and the total number of variants *n* identified in the region [pbinom(q, n, p, lower.tail = FALSE)].

### Coding Variant Panel Design

To profile the coding genome in these patients, a refined panel of genes known as the Oncomine panel was utilised, specifically designed to cover key areas of mutation in luminal breast cancers^78^. The panel targets 6,812 coding regions, selected by compiling commonly mutated sites identified in up-to-date studies, sequencing both primary and metastatic luminal breast cancer tumours. The panel utilised data from an array of databases and studies including: The Cancer Genome Atlas (TCGA) database, the Molecular Taxonomy of Breast Cancer International Consortium (METABRIC) database ^79^, Lefebvre et al 2016 ^80^, the MSKCC IMPACTTM study ^81^, the AACR GENIE database ^82^, the COSMIC database, the Cancer Gene Census, and the Pharmacogenomics Knowledgebase (PharmKGB)^83^. In total, these datasets included 1,673 primary and 1,596 metastatic luminal breast cancer cases. Mutated genes identified in these datasets were compiled and refined using the following criteria. Sites that were mutated in at least 2% of primary or metastatic samples and CNVs with a frequency of over 5% or with a fold change of over 5% in either primary or metastatic tumours were compiled. All breast cancer genes reported in the Cancer Gene Census and all pharmacogenomic SNPs related to breast cancer in the PharmKGB database were compiled. Finally, some manual curation was included, adding in the CYP19A1 and SQLE amplification^9, 84^. After refinement, the panel included 6,812 regions covering 134 genes, 27 CNV sites, 37 germline cancer genes, and 59 germline loci, with associations to pharmacogenomic interactions.

### Sample preparation and sequencing

Secondary captures, on SIDV, captured DNA libraries, was carried out using the Oncomine panel. After hybridisation of SIDV capture probes to complementary DNA and purification, non-captured DNA was recovered and concentrated using SPRI size-selection beads. Quality control assessment using a Tapestation 2200 instrument was performed reporting that, in all cases, at least 50% recovery of initial DNA concentrations before the SIDV capture had been achieved. A custom set of capture probes for the Oncomine regions were produced by Twist Biosciences. Pools of DNA were captured using the Oncomine panel and quality controlled as previously described with the SIDV panel. Pools of 10 patients were sequenced at Novogene on an Illumina NovaSeq 6000 (150bp paired-end), with 700 million reads per pool.

#### Computational analysis of Coding Variants

Variant calling was initially performed for all 100 patients that were sequenced – matched normal, primary and metastatic samples. Adapter trimming was performed using Trim Galore version 0.6.4 (https://www.bioinformatics.babraham.ac.uk/projects/trim_galore/). Bwa-mem version 0.7.15 PMID: 19451168 was used for alignment to the hg38 human genome reference. Sambamba ^85^version 0.7.0 was used for conversion to binary, removal of PCR duplicates, sorting and indexing. Pre-processing before variant calling was performed using GATK^86^, version 4.1.3.0: read groups were added using picard version 2.20.6 (https://sourceforge.net/projects/picard/files/picard-tools/), base quality recalibration using gatk BaseRecalibrator and gatk ApplyBQSR. Mutect2 was used for somatic variant calling against the matched normal bam samples: using the germline resource from the GATK resource bundle af-only-gnomad.hg38.vcf.gz with option –af-of-alleles-not-in-resource set as 0.001 and with MateOnSameContigOrNoMappedMateReadFilter disabled. To flag possible FFPE artefacts gatk LearnReadOrientationModel was run, using output during the filtering of variants with FilterMutectCalls. Only PASS mutations were further processed. Depth was checked at 500 mutated loci (variants with a FATHMM score >= 0.8 and a variant allele frequency (VAF) of at least 0.1 from the pool of de novo metastatic mutations) in all 100 patients – across normal, primary and metastatic - using samtools depth. This analysis revealed that in 42/100 patients, depth was lower than 10 in the majority of the loci, in at least one of the normal, primary or metastatic bam files. Since this low number of reads could affect variant detection generally, or affect the identification of de novo metastatic variants (i.e. impossible to discern whether a mutation found in the metastatic sample was not present in the primary if the depth at that locus is low in the primary). As depth was sufficient across all variants in the other 58 patients, these were further processed. Variant annotation was performed using OpenCRAVAT, filtering for mutations only found in established breast cancer driver genes^87^. To discover potential de novo driver variants of metastasis in these patients, we filtered for non-synonymous coding variants, with >= 0.1 VAF, private to metastasis or with an allele frequency at least 5 times higher than in the primary. ComplexHeatmap version 2.9.3. (http://bioconductor.org/packages/release/bioc/html/ComplexHeatmap.html) was used to generate an OncoPrint heatmap of these de novo, possibly pathogenic variants.

#### CRISPRi screen: sgRNA design

First, promoter-associated SIDV3 regions were excluded (a more tailored design of sgRNAs guided by available CAGE tags data in MCF7 was performed instead, see below for details). After enlarging each region to be at least 500 bps in size, the command-line version of the CRISPR-DO tool (version 0.04,^88^) was then run separately for each one of the considered regions (with --spacer-len=20), and the predicted sgRNAs stored. Only sgRNAs showing efficiency between 0.4 and 1.3, and specificity >= 80% were retained for further analyses. One G nucleotide was then added at both 5’ and 3’ of each sgRNA, and the resulting guides predicted to be digested by endonuclease BbsI were discarded. In silico digestion was performed using the *digest* package in R. After that, to obtain a more uniform distribution of sgRNAs, an iterative pruning procedure was applied until no two guides were found within 50 bps from each other. This resulted in 62.2% and 79.7% of the putative insulators and enhancers showing 3 or more sgRNAs targeting them, respectively. Only the sgRNAs targeting those regions were retained.

Hg19 coordinates for CAGE tags peaks from FANTOM5 ^89^ were downloaded from the consortium website (https://fantom.gsc.riken.jp/5/datafiles/latest/extra/CAGE_peaks/). Briefly, starting from hg19.cage_peak_phase1and2combined_tpm_ann.osc.txt.gz, only those expressed at least with a TPM >= 1 in unstimulated MCF7 were considered further. For each gene (after filtering for blacklisted regions in ENCODE and for promoters of anti-sense, non-coding RNAs) the dominant TSS (based on highest CAGE TPM) was identified. Only a single, dominant TSS for each expressed gene was retained. Of those, only those corresponding to promoters of genes with at least one overlapping putative insulator or enhancer in SIDV3 were considered for sgRNA design. Considering the directionality of transcription at each CAGE tags cluster, each region was standardized to [-100, +300] bps from the dominant position in the cluster. Design and filtering of the sgRNAs were then performed as described in the previous paragraph.

#### CRISPRi screen: data analysis

Count data were normalised according to the weighted trimmed mean of the log expression ratios (trimmed mean of M values (TMM)) normalisation^90^, using the *calcNormFactors* function from edgeR^91^. Initial PCA and clustering analyses indicated high similarity between the 8 days samples and the initial library. For this reason, the replicated 8 days samples were used as a reference to identify statistically significant changes in abundance of sgRNAs at later time points, using edgeR^91^. Briefly, after estimating dispersion using the *estimateDisp* function, generalised linear models (GLMs) were fit separately to each condition (full and oestrogen-depleted medium), using the *glmFit* function. Coefficients were retrieved with *glmLRT*, and significant changes were retained as those showing a Benjamini-Hochberg corrected FDR <= 0.05 and a log2-fold-change of at least 1, in either direction. The same computational strategy was applied to compare the sgRNAs counts in full vs oestrogen-depleted media, at any given time point.

### Statistical analyses and plotting using R

Unless indicated otherwise, all the described statistical analyses and preparation of plots were performed in the statistical computing environment R v4 (www.r-project.org).

## Supplementary Figures Legends

**Supplementary Figure 1.**
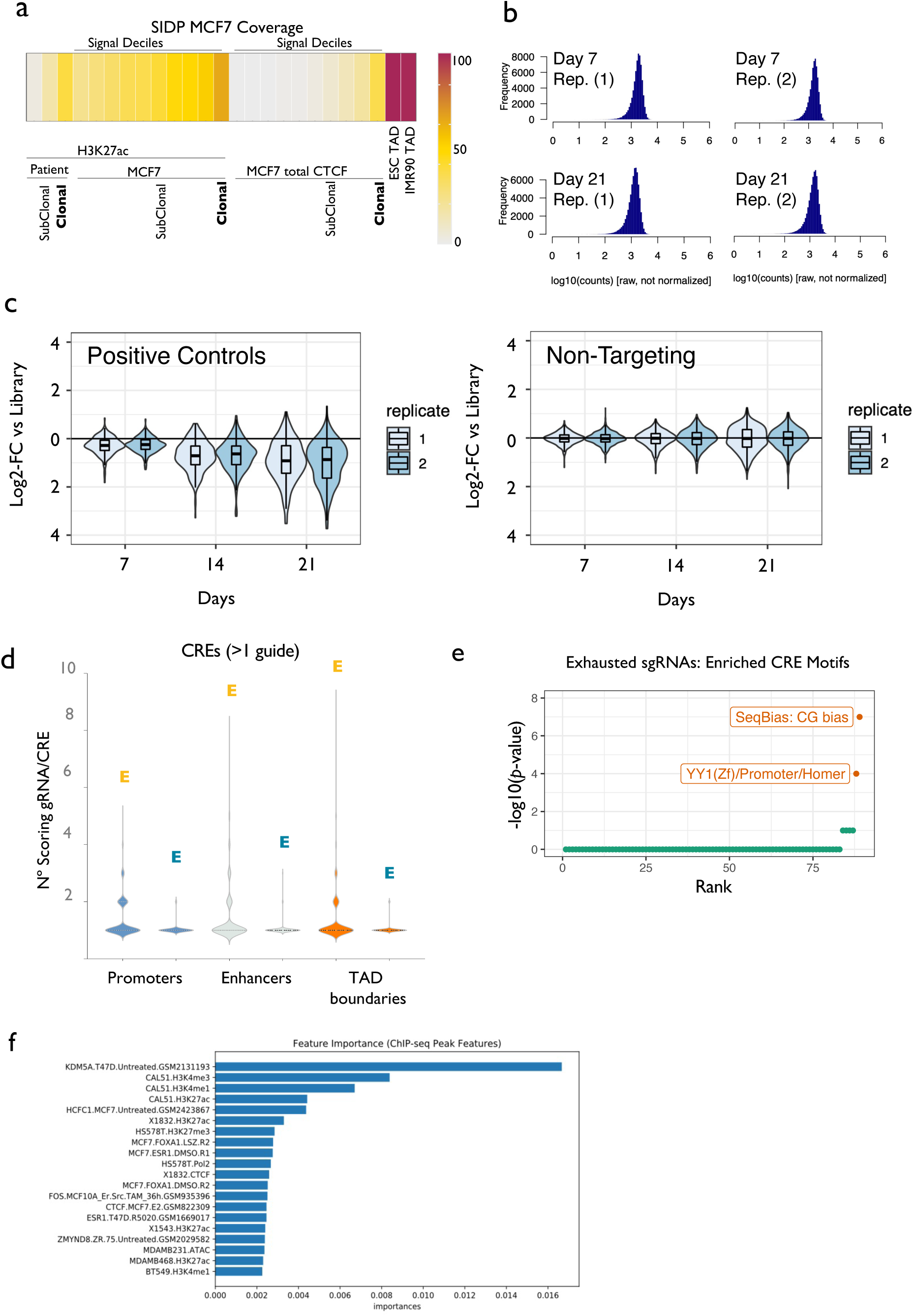
**(a)** SIDP coverage (percentage) of the specific partitions of the human CREs considered in this study. **(b)** Histograms showing the distribution of counts per sgRNAs (log10) for two replicates of sgRNAs in pool 1, at day 7 and day 21 post-infection (MCF7 full media). **(c)** Box plots showing the log2-fold-change of positive controls (left panel) and non-targeting sgRNAs (right panel) in two replicates of oestrogen-dependent MCF7 cells, at 7, 14 and 21 days, as compared to the initial library. **(d)** Box plots showing the distribution of the number of significantly scoring sgRNAs per CRE, for Expanded (yellow) and Exhausted (blue) sgRNAs, across three different genomic partitions (promoters, putative enhancers, and CTCF-clusters associated to TAD boundaries). **(e)** Motif analysis of CREs associated with significantly exhausted sgRNAs identifies YY1 as a putative TF enriched in functional CREs.

**Supplementary Figure 2.**
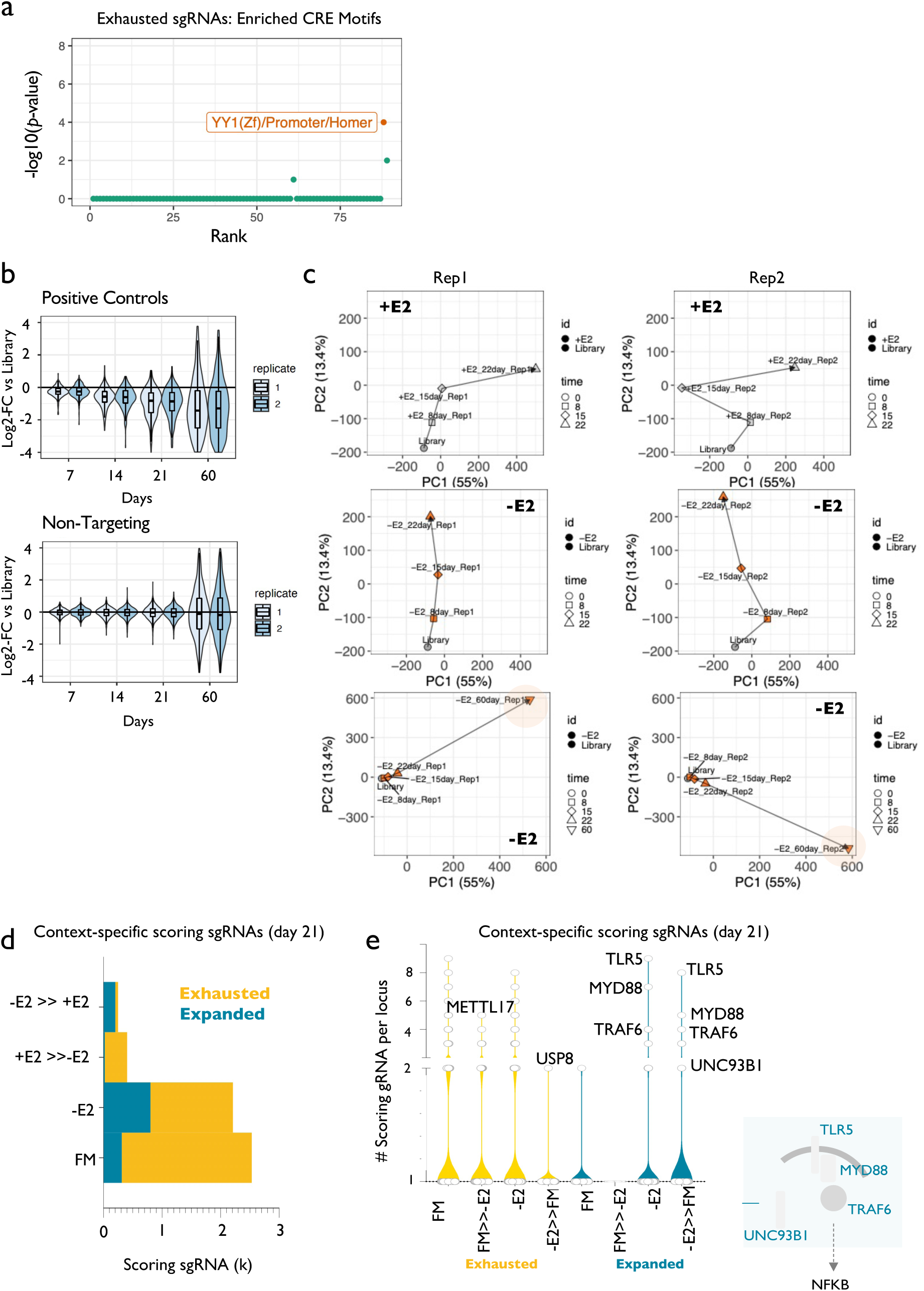
**(a)** Motif analysis of CREs associated with significantly exhausted sgRNAs identifies YY1 as a putative TF enriched in functional CREs. **(b)** Box plots showing the log2-fold-change of positive controls (left panel) and non-targeting sgRNAs (right panel) in two replicates of oestrogen-deprived MCF7 cells, at 7, 14 and 21 days, as compared to the initial library. **(c)** Principal component analysis (PCA) of all samples. For clarity in the visualization, the analysis was first run excluding day 60 and split by replicate (columns) and condition (+/- E2; first two rows). The last pair of plots from above, instead include day 60. Note: 8 and 15 days post cells seeding (corresponding to 7 and 14 post-infection). **(d)** Bar plot showing the overall number of sgRNAs showing the indicated behaviour ad day 21 (MCF7 white media). **(e)** Box plots showing the number of sgRNAs significantly decreased or increased at day 21 (MCF7 white media). Specific outliers indicate the nearest gene to the overlapping CRE. (Bottom right) Schematic of the genes identified in the TLR/NF-kB signalling pathway, showing at least one CRE with multiple expanded sgRNAs at day 21 (MCF7 white media).

**Supplementary Figure 3.**
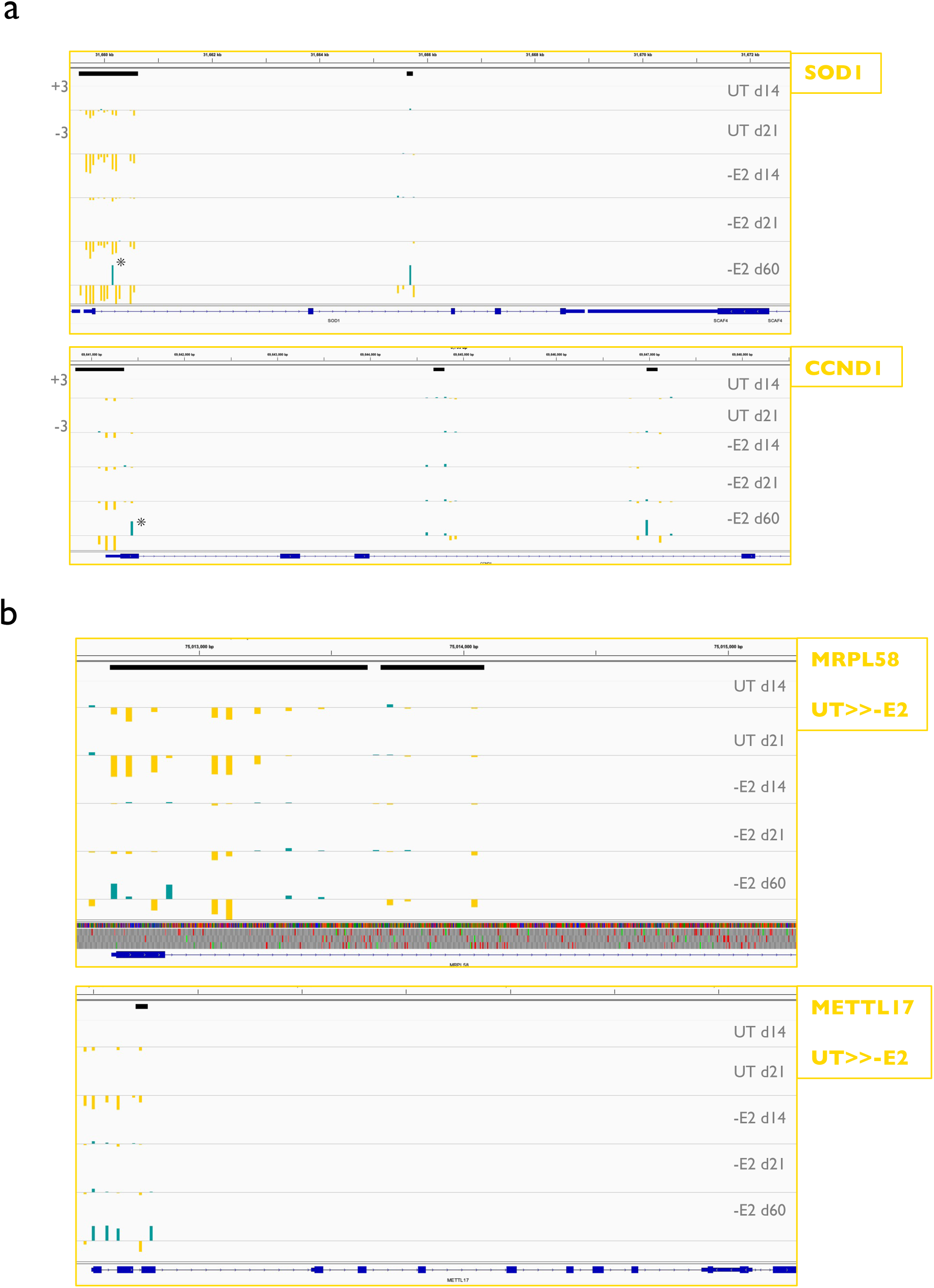
**(a-b)** SIDP results at the indicated loci are shown as IGV genome browser screenshots. For each of the indicated conditions, the log2-fold change for each sgRNA is indicated, with bars proportional to the effect size, and colour reflecting the sign (blue = expanded; yellow = exhausted).

**Supplementary Figure 4.**
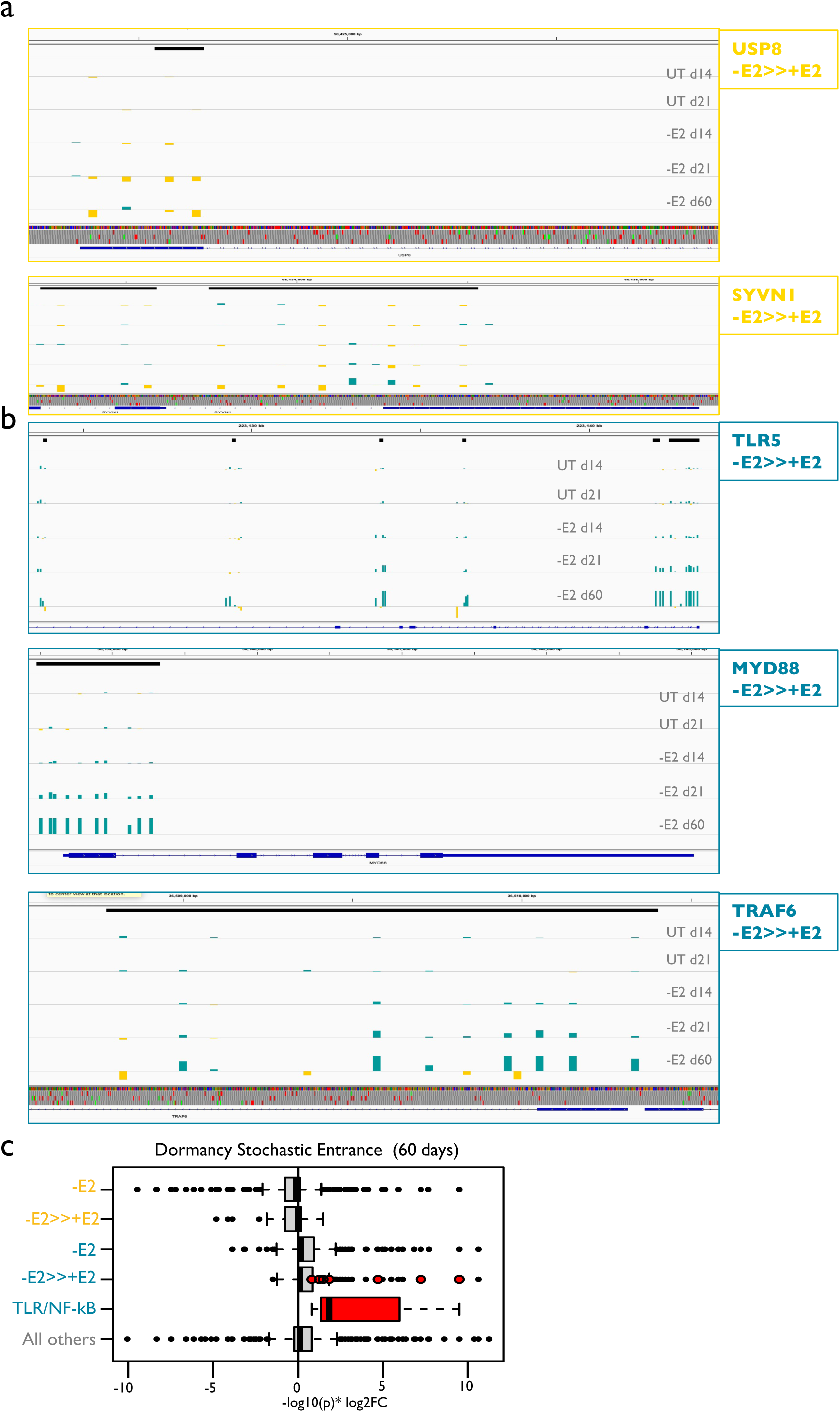
**(a-b)** SIDP results at the indicated loci are shown as IGV genome browser screenshots. For each of the indicated conditions, the log2-fold change for each sgRNA is indicated, with bars proportional to the effect size, and colour reflecting the sign (blue = expanded; yellow = exhausted). **(c)** Box plots showing the distribution of the compounded score (-log10 of the edgeR-estimated *p*-value times the log2FC) for different sets of sgRNAs (blue = expanded; yellow = exhausted), at 60 days (MCF7 white media). The scoring sgRNAs mapping to the CREs of the genes annotated to TLR/NF-kB signalling are highlighted in red (as outliers in the distribution of the sgRNAs significantly more expanded in -E2 vs +E2 conditions at 21 days, and then as a separate group).

**Supplementary Figure 5.**
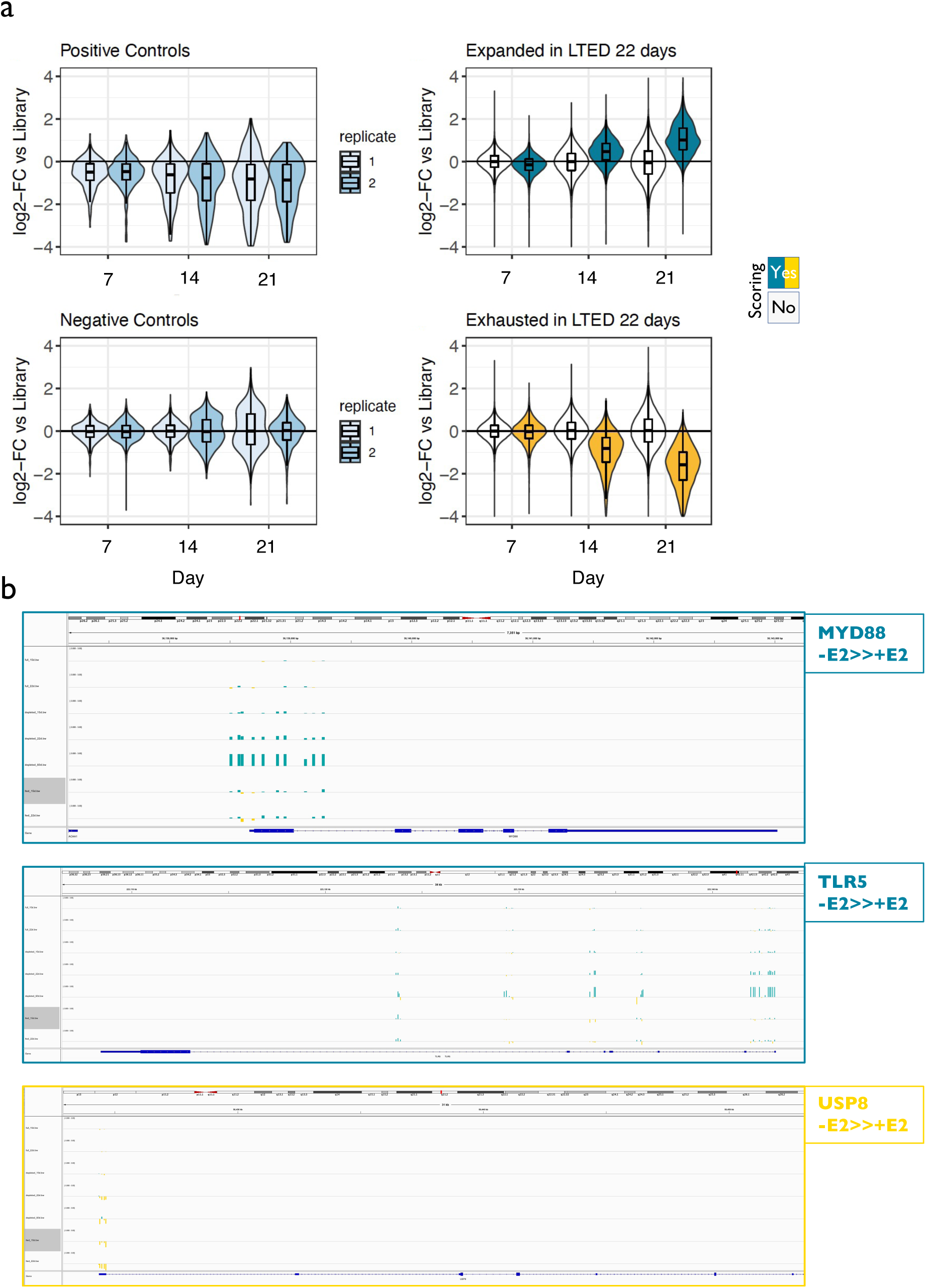
**(a)** (top to bottom, left to right) Box plots showing the log2-fold-change of positive controls and non-targeting sgRNAs in two replicates of oestrogen-deprived MCF7 cells, at 7, 14 and 21 days, as compared to the initial library. The other two box plots show log2-fold-change of both scoring (either blue or yellow) and non-scoring (white) sgRNAs at 21 days post-infection in oestrogen-deprived MCF7 cells, at 7, 14 and 21 days, as compared to the initial library. **(b)** SIDP results at the indicated loci are shown as IGV genome browser screenshots. For each of the indicated conditions, the log2-fold change for each sgRNA is indicated, with bars proportional to the effect size, and colour reflecting the sign (blue = expanded; yellow = exhausted).

**Supplementary Figure 6.**
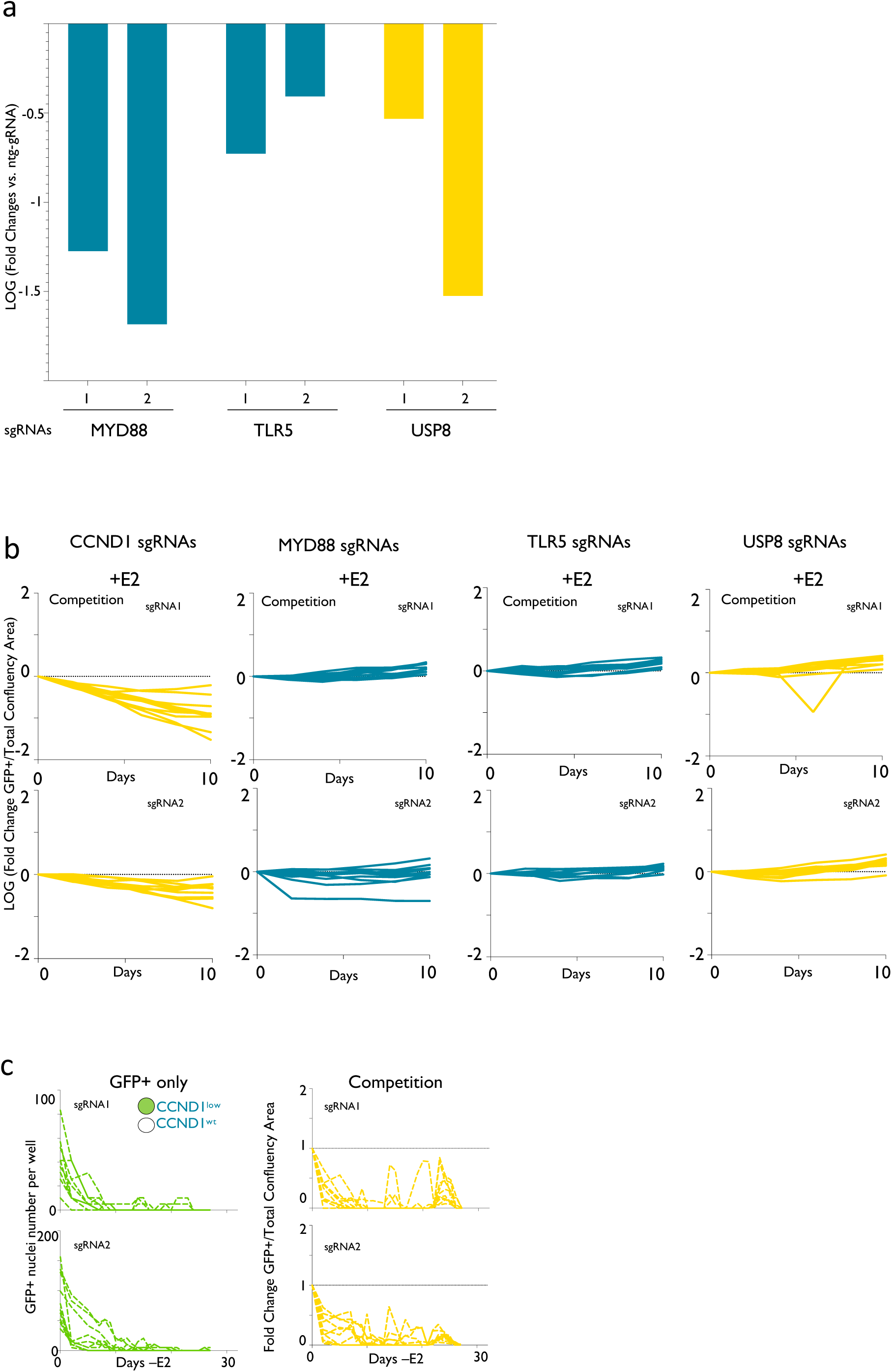
**(a)** RT-qPCR validation of the effect of individual sgRNA on MCF7 transfected with a constitutive dCAS(-KRAB construct. Relative mRNA values are plotted against a non-targeting sgRNA **b)** Cell competition experiments. 150 cells GFP positive transfected with single experimental sgRNAs were plated with 150 cells transfected with non-targeting sgRNA. The relative ratio of GFP+/non-GFP cells across ten days is plotted. Experiments were conducted in full media (with estradiol) **c)** CCND1 targeting sgRNAs lead to the rapid extinction of GFP cells while non-targeted cells enter dormancy with normal dynamics. Green panels: absolute GFP+ count (CCND1 sgRNAs). Yellow panels: normalized ratios GFP/non GFP across time points.

**Supplementary Figure 7.**
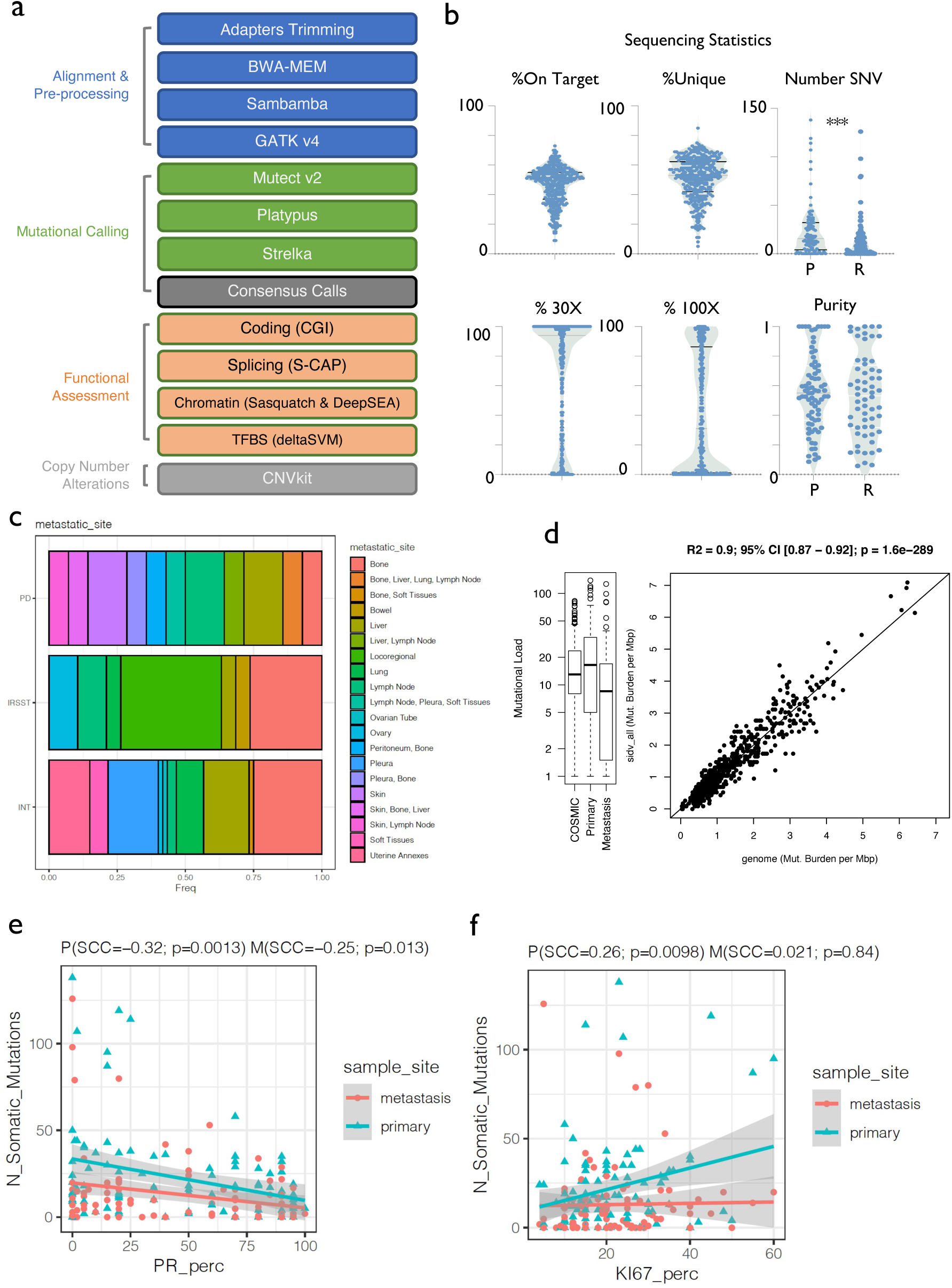
**(a)** Schematic summarising the steps of the custom computational pipeline employed for the identification and functional annotation of the SIDV variants. **(b)** Summary of the sequencing statistics for the profiled samples. **(c)** Stacked bar plots showing the anatomic site of the profiled relapse, split by centre. **(d)** Box plot showing the distribution of the overall mutational load per sample in the SID regions, as estimated in either the non-coding COSMIC or in our cohort (separately for primary and metastatic samples). The companion scatterplot shows the correlation between the genome-wide estimate of mutational burden and the same estimate using only the mutations identified in SID regions, considering the WGS data available in the non-coding COSMIC. The statistics and statistical significance of this linear correlation are indicated on top of the plot. **(e-f)** Scatterplots showing the relationship between the number of somatic mutations detected per sample, and the indicated variables. For visualization purposes, least-square regression models were trained separately for primary and metastatic samples. For quantifying the relationships, Spearman’s correlation coefficients (SCC) are indicated on top of the plots, along with the corresponding *p*-values.

**Supplementary Figure 8.**
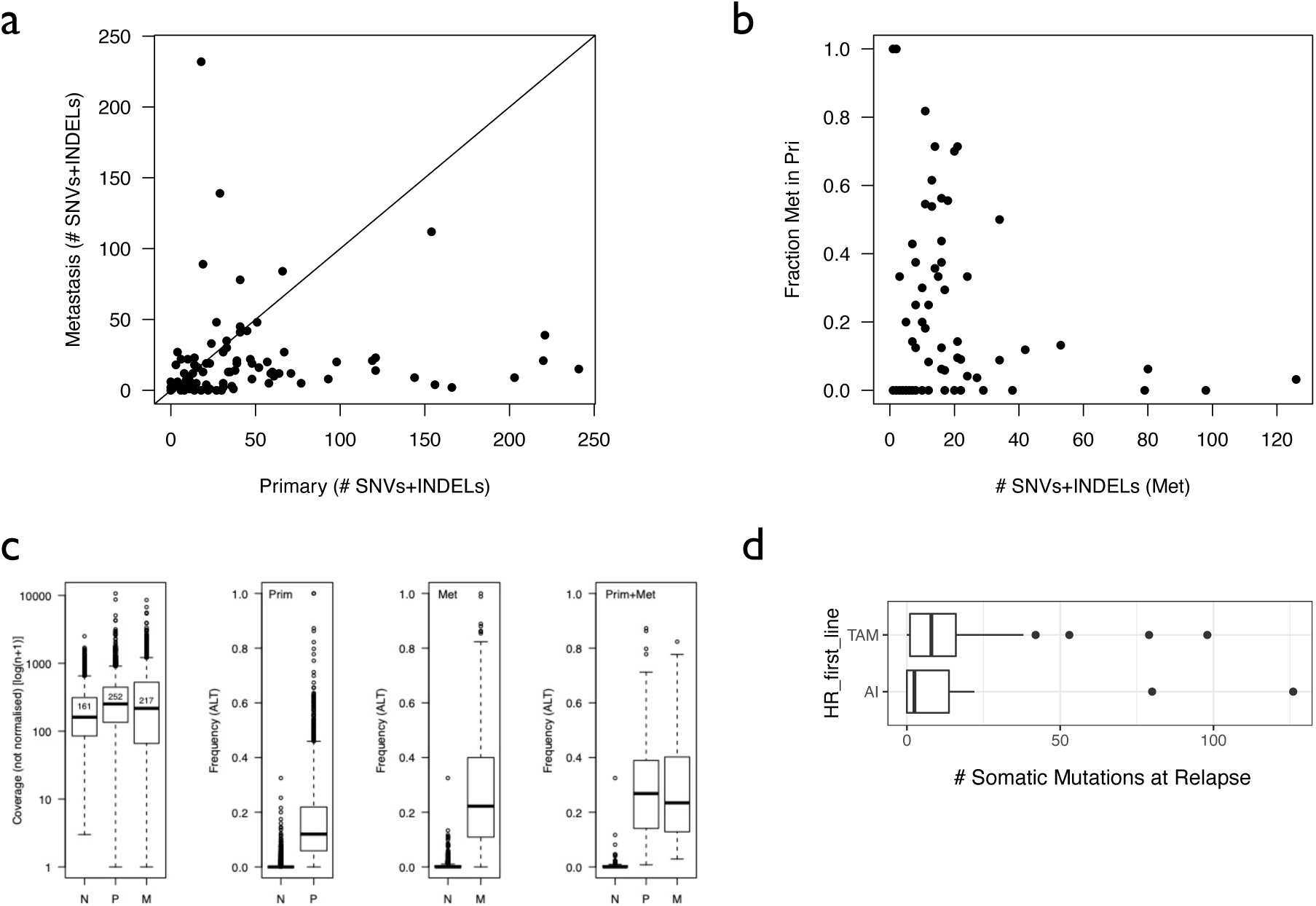
**(a)** Scatterplot comparing the number of variants (SNVs plus INDELs) in matched primary and metastatic lesions. **(b)** Scatterplot showing the fraction (0-1) of mutations identified in the metastatic tumour that was also called in the corresponding matched primary. Each dot represents a pair of matched primary-met, with the x-axis indicating the total number of variants in each metastatic sample. **(c)** (Left to right) Box plots showing the overall coverage of the regions showing variants, separately for matched normal (N), primary (P) and metastatic (M) samples. The other three box plots show the VAF (frequency of alternative alleles) in normal and tumour (either primary, metastatic, or both) specimens, for three sets of variants (left to right): those identified only in primaries; those identified only in metastasis; those identified in both. **(d)** Box plot showing that lesions that were treated with TAM or AI did not show a different number of detected mutations at relapse (*p*-value = 0.21; Mann-Whitney Test).

## Supplementary Tables Legends

**Supplementary Table 1: Regions defined by SID (Systematic Identification of epigenetically Defined loci).** For each region (hg38 genomic coordinates including chromosome, starting and ending positions), the table indicates whether the region was selected as a gene promoter, putative enhancer or putative insulator. Whether the region is covered by designed oligo baits for SIDV profiling, and the number of sgRNAs targeting the region in SIDP, are also indicated.

**Supplementary Table 2: sgRNAs sequences and metadata for the SIDP assays. S2.1:** for each sgRNA targeting a region in the human genome, an identifier (using the corresponding hg38 genomic coordinates), the DNA sequence, the genomic coordinates (hg38) including the strand, along with efficiency and specificity scores as estimated by CRISPR-do, are provided. **S2.2:** for each positive control or non-targeting sgRNA, a custom identifier is shown along with the DNA sequence.

**Supplementary Table 3: SIDP results in MCF7 grown in full (red; +E2) media. S3.1:** results of the differential abundance analysis for the positive controls and the non-targeting sgRNAs (as indicated in the genome_partition field). For each sgRNA, an identifier, the pool, and the results from the edgeR analysis are shown. The average abundance of the sgRNA at day 7 and 21 post-infection is indicated as logCPM (counts per million). The log2-fold changes (log2FC) between day 21 and 7, and between day 21 and the initial library, are indicated, along with the FDR (Benjamini-corrected *p*-value). Two further fields indicate whether the sgRNA was identified as significantly expanded (FDR <= 0.05 and linear fold-change >= 1.5) or exhausted (FDR <= 0.05 and linear fold-change <= -1.5). **S3.2:** similar to S3.1 but listing the results for the sgRNAs targeting the genomic regions of interest. Hg38 coordinates are also included in this case. **S3.3:** summary of the results at the level of each SID region. For each region, hg38 coordinates are listed, along with the symbol of the nearest gene, and the distance to its TSS in bp (positive or negative values indicate the region is either downstream or upstream the TSS, respectively). The table then indicates whether the region was selected as a gene promoter, putative enhancer or putative insulator. The number of sgRNAs targeting the enlarged region (indicated coordinates +- 1 kbp), is followed by information on the overlapping sgRNAs that scored significantly, separately for exhaustion and expansion. In both cases, the total number of significant guides, the corresponding fraction, and the FDR and log2FC of the highest-scoring sgRNA are reported. A column indicating the significance of one or more sgRNAs is also provided. **S3.4:** enriched terms in the set of genes close to the regions showing scoring sgRNAs, separately for the exhausted and the expanded sets. For each group, hallmark sets showing a *p*-value <= 0.05 are included in the table. Statistics of the hypergeometric test are shown, along with the total number and identity of the overlapping genes. **S3.5:** overlap between the regions identified in our +E2 MCF7 SIDP assay and previously published screens in breast cancer cell lines (marcotte: Marcotte et al. 2012; fei: Fei at al. 2019; Korkmaz: Korkmaz et al. 2019; ggg: Rui Lopes et al. 2020).

**Supplementary Table 4: SIDP results in MCF7 grown in white media (-E2). S4.1-5:** the tables follow the same structure as S3.1-5.

**Supplementary Table 5: SIDP results in LTED. S5.1:** results of the differential abundance analysis for the positive controls and the non-targeting sgRNAs. The structure of the table is similar to S3.1. **S5.2:** results for the sgRNAs targeting the genomic regions of interest. The structure of the table is similar to S3.2.

**Supplementary Table 6: SIDP results summary. S6.1:** regions showing at least one overlapping sgRNA scoring in at least one of the different conditions assayed. For each region (hg38 genomic coordinates), the table indicates whether this was selected as a gene promoter, putative enhancer or putative insulator. It also shows the symbol of the nearest gene, and the distance to its TSS in bp (positive or negative values indicate the region is either downstream or upstream of the TSS, respectively). For each condition (MCF7 RM, MCF7 WM or LTED) and direction of the change (Exhaustion vs Expansion), the table indicates whether the region overlaps one or more (columns labelled “single”) vs two or more (columns labelled “multiple”) sgRNAs. **S6.2:** summary of the overlaps between either scoring sgRNAs (“guides”), regions showing at least one scoring sgRNA (“regions_single”), or regions showing two or more consistently scoring sgRNAs (“regions_multiple”) between pairs of conditions (as indicated by columns assay_1 and assay_2). The nature of the change (either Exhaustion or Expansion), along with the total number of overlapping sgRNAs or regions, and the corresponding fraction, are also indicated. **S6.3:** results of gene set enrichment analysis using the indicated gene sets and the set of genes close to the regions showing scoring sgRNAs, according to the indicated pattern (SIDP_set). Statistics of the hypergeometric test are shown, along with the total number of the overlapping genes (count), the observed and expected overlaps, and the odds ratio.

**Supplementary Table 7: Metadata of the clinical cohort profiled by SIDV. S7.1:** for each donor, from which genetic material from matched normal, primary and metastatic samples was derived, the following information is provided: the identifier for the samples; the centre where the samples were collected; the sequencing batch; the age of diagnosis; the clinical features of the primary tumours; the indication of the metastatic sites. Legend: ER = estrogen-receptor alpha; PR = progesterone receptor; pct = percentage; HR = hormone therapy. **S7.2:** for each triplet of matched normal, primary and metastasis derived material, and separately for each one of the 100 donors, sequencing statistics are provided. Sequencing depth, the fraction of the reads mapping to oligo baits, mean coverage on baits and corresponding fold-enrichment, and on-target mean coverage, are shown. The percentages (pct) of targeted bases covered at least 10x, 30x, 50x or 100x are also indicated.

**Supplementary Table 8: Summary of SNVs and INDELs identified by SIDV. S8.1:** total number of SNVs and INDELs (filtered for common variants, according to dbSNP) per donor (sample_id), divided by those identified in primary or metastasis (vs matched normal). **S8.2:** full list of SNVs and INDELs. Chromosome and position on the chromosome (hg38 coordinates) are indicated for each variant, along with the reference and detected alternative allele. Also, the table indicates the donor, and whether the variant allele was directly detected in the primary (P_CALL) and/or the metastatic material (M_CALL). **S8.3:** tumour purity estimation for each sample and site (P = primary; M = metastasis) is listed, along with the size of the subset of SNVs used for the purity estimation analysis. **S8.4:** final annotation of the SNVs after sample-specific purity correction. For each SNV, genomic coordinates, reference and alternative alleles, donor identifier, and evidence (filtered read counts) supporting the different alleles in normal (N), primary (P) and metastatic (M) samples are provided. For both primary and metastatic samples, the variant allele frequency (VAF), along with the estimated purity for the sample, the estimated copy number alterations of the region bearing the variant (CNA) and the purity-corrected VAF, or cancer-cell fraction (CCF), are indicated. **S8.5:** regions showing an enrichment in either amplification (amp) or deletions (del) across the metastatic samples as compared to the matched primary samples, are indicated.

**Supplementary Table 9: Computational predictions of the functional impact of the SNVs and short INDELs identified through SIDV. S9.1:** for each variant, the type (SNV or INDEL_short) and its hg38 coordinates are listed, along with the symbol of the nearest gene, and the distance to its TSS in bp (positive or negative values indicate the region is either downstream or upstream the TSS, respectively). Reference and alternative alleles are also provided, along with whether the variant is computationally predicted to alter the molecular function of the genomic element bearing it (indicated as different “pathogenic” classes; column mutation_class) or not (“benign”). The table is then indicating, for each one of the models considered, whether the variant is predicted to significantly affect the indicated molecular function. **S9.2:** extract of S9.1, for three regions of interest.

**Supplementary Table 10: Downstream analyses considering only the SIDV inferred genetic alterations with predicted impact on function. S10.1:** results of the binomial enrichment test. SID regions overlapping at least 2 SNVs predicted as pathogenic are included. Along with genomic coordinates (hg38) the total number of SNVs, as well as the number of predicted pathogenic SNVs overlapping the region, are indicated. The *p*-value and the *q*-value (after Benjamini-Hochberg correction) of the binomial test are indicated, along with annotation to the closest gene. **S10.2:** same as S10.1, but considering all the regions assigned to the genes annotated to the same ontological terms together. The number and identity of the genes contributing to the overlap are indicated, along with the *p*-value of the binomial test, and the *q*-value (after Benjamini-Hochberg correction). Statistically significant terms (*q*-value <= 0.05) are highlighted in red. **S10.3:** results of the analyses testing for the enrichment of mutations (either SNVs, short INDELs, or both; mutation_type column) with computationally predicted pathogenic effects in the sets of regions also showing a certain behaviour in SIDP (CRISPRi_hit_type column). Observed and expected overlap are indicated, along with the odds ratio and the *p*-value (Chi-squared test).

**Supplementary Table 11: Downstream analyses considering only the SIDV inferred genetic alterations with predicted impact on function, and stratifying them by cancer-cell fraction (CCF) increase and decrease in metastatic samples. S11.1:** summary of the results of the statistical tests performed to identify differences in the predicted impact of mutations stratified by a change in CCF in metastatic samples compared to matched primary. The fraction of variants predicted as pathogenic and either showing an increase or a decrease in CCF (+- 0.1) was compared to that of those showing no change. *P*-values for the indicated features are shown (Chi-squared test). **S11.2:** similarly, the distribution of the predicted molecular effects of variants in the three groups (increase, decrease or no change in CCF) were compared using the Kruskal-Wallis test. **S11.3:** similar to S10.3, but testing for the enrichment of mutations with both computationally predicted pathogenic effects and a certain CCF increase or decrease in metastatic samples, that also show a certain behaviour in SIDP.

**Supplementary Table 12: Results of the enrichment analyses looking for binding sites of specific TFs accumulating more or less genetic variants than expected by chance.** For each TF and category (mutations significantly increasing or decreasing affinity) the observed and expected fraction of mutations overlapping the TF-bound sites are indicated, along with the difference between these two fractions, and the *p*-value of the corresponding Chi-squared test. Considering each TF and the mutations affecting the affinity to its target sites either positively or negatively (based on the *p*-value of the test) TFs could be either classified as showing significantly more or fewer mutations than expected, or not significant (ns).

**Supplementary Table 13: Datasets used for the training of the TF-specific deltaSVM models.** For each TF, the corresponding gene symbol, along with information about the cells from which the ChIP-seq binding profile was obtained, the treatment the cells were exposed to (if any), and reference to the corresponding records on the Gene Expression Omnibus, are indicated. Information about the matched, high-quality position weight matrix (PWM) utilized as a source of information to infer the binding affinities of each TF is also provided. For each PWM, an identifier is indicated, along with the corresponding reference database or publication (including Pubmed ID).

